# High-density volumetric super-resolution microscopy

**DOI:** 10.1101/2023.05.02.539032

**Authors:** Sam Daly, João Ferreira Fernandes, Ezra Bruggeman, Anoushka Handa, Ruby Peters, Sarah Benaissa, Boya Zhang, Joseph S. Beckwith, Edward W. Sanders, Ruth R. Sims, David Klenerman, Simon J. Davis, Kevin O’Holleran, Steven F. Lee

## Abstract

Volumetric super-resolution microscopy typically encodes the 3D position of single-molecule fluorescence into a 2D image by changing the shape of the point spread function (PSF) as a function of depth. However, the resulting large and complex PSF spatial footprints reduce temporal resolution by requiring lower labelling densities to avoid overlapping fluorescent signals. We quantitatively compare the density dependence of single-molecule light field microscopy (SMLFM) to other 3D PSFs (astigmatism, double helix and tetrapod) showing that SMFLM enables an order-of-magnitude speed improvement compared to the double helix PSF by resolving overlapping emitters through parallax. We then experimentally demonstrate the high accuracy (>99.2 ± 0.1%, 0.1 locs μm^−2^) and sensitivity (>86.6 ± 0.9%, 0.1 locs μm^−2^) of SMLFM at point detection through whole-cell (scan-free) imaging and tracking of single membrane proteins in live primary B cells. We also exemplify high density volumetric imaging (0.15 locs μm^−2^) in dense cytosolic tubulin datasets.

---

Single-molecule localization microscopy (SMLM) is a super-resolution technique that separates the fluorescence emission of individual fluorophores temporally to observe biological systems with sub-diffraction resolution (1–4). Direct imaging in three dimensions (3D) enables the study of complex biological morphologies and dynamic processes that would otherwise be underestimated in 2D.

In SMLM, the fluorescence from a single emitter is observed as a diffraction-limited spot on a detector, known as the *point spread function* (PSF). Generally, 3D-SMLM employs optical elements that transform the standard 2D PSF into spatial distributions that also encode axial position. These 3D PSFs exhibit lateral spatial footprints that are much larger in area than the standard PSF, meaning the projection of a 3D volume onto a 2D detector usually necessitates considerably slower acquisition rates (typically 5 to 10-fold) due to a higher likelihood of PSF overlap (5, 6). However, the number of emitters localised per frame governs temporal resolution and therefore dense emitter datasets are desirable. This is exemplified in recent work from Legant *et al*. where impressive super-resolved whole-cell volumes were obtained over very long acquisition times (*i*.*e*. 3–10 days) (7, 8). This extended experimental duration was necessary to generate an image with resolution comparable to a corresponding electron microscope experiment (8).

Long imaging durations present unrealistic conditions for typical cellular experiments and also reduce the quantity of biological repeats that can be performed within appropriate timescales. While strategies exist to reduce PSF overlap—such as specialised labelling protocols and post-processing algorithms (9–11)—they are ultimately limited by the decrease in lateral resolution at the expense of a greater depth-of-field (DoF). However, a recent study revealed a lack of post-processing solutions specifically for dense 3D datasets (12). Hence, to have broad applicability to the biological community there is a fundamental need for robust strategies to perform 3D-SMLM at high densities. This will bring 3D-SMLM into line with the timescales and workflows of current 2D cellular experiments, and is another important step toward real-time 3D-SMLM.

Sub-diffraction axial precision can be achieved by engineering the shape of the PSF to simultaneously encode the lateral and axial position of a single emitter in a 2D image (13, 14). A variety of engineered PSFs have been reported, including astigmatism (∼1 μm DoF) (15), a bisected pupil (∼1 μm DoF) (16), the corkscrew PSF (∼3 μm DoF) (17), the double helix (DH)PSF (∼4 μm DoF) (5, 18, 19), and the tetrapod PSF (6–20 μm DoF) (20, 21). On the other hand, single-molecule light field microscopy (SMLFM) (22) is an SMLM technique that places a refractive microlens array (MLA) in the back focal plane (BFP) of a widefield microscope to encode 3D position into the PSF (see Supplementary Note S1) (23, 24). SMLFM possesses a large tuneable DoF, high photon throughput, (13) the PSF can be fitted with conventional 2D algorithms and is wavelength non-specific. The unique advantage of SMLFM is that it operates through parallax whereby the PSF is comprised of several spatiotemporally correlated perspec-tive views displaced in proportion to the curvature of the wavefront. As such, SMLFM is particularly suited to high spot densities for two key reasons:

1. Single emitters that occur at different axial planes (but overlap laterally) are imaged at different locations in different perspective views and can be distinguished.
2. We illustrate a redundancy in that a localisation is not required in every perspective view to be localised in 3D.

Multi-focal plane microscopy also segments the BFP to image two or more focal planes and capture 3D volumes (25–28). However, this work will focus on techniques that yield sub-diffraction axial precision over extended axial ranges.

In the present work, we report the first hexagonal SMLFM platform capable of super-resolving single emitters at very high densities over an 8 μm DoF. We quantitatively compare the performance of SMLFM to other common 3D PSFs as a function of spot density through simulations. We then apply SMLFM experimentally to the scan-free imaging and tracking of individual B-cell receptors and the imaging of tubulin in HeLa cells to show that overlapping PSFs minimally affect the localization precision and that existing labelling strategies can now be directly transferred to 3D imaging pipelines.

## Results and Discussion

### Density dependence of 3D PSFs

Current state-of-the-art 3D PSFs are typically created by phase modulation in the BFP of the objective lens (*i*.*e*. with a phase mask) as shown in Fig. 1a. This phase modulation gives rise to spatial distributions of intensity in the imaging plane that change as a function of the axial position of the emitter. Alternatively, in SMLFM the MLA segments the BFP and focuses an array of spots on the detector as shown in Fig. 1b and Supplementary Fig. S1. All of these changing PSFs can be understood in the context of high-density imaging by collapsing the entire PSF onto the 2D detector, which we define as the *PSF footprint*.

**Fig. 1.**
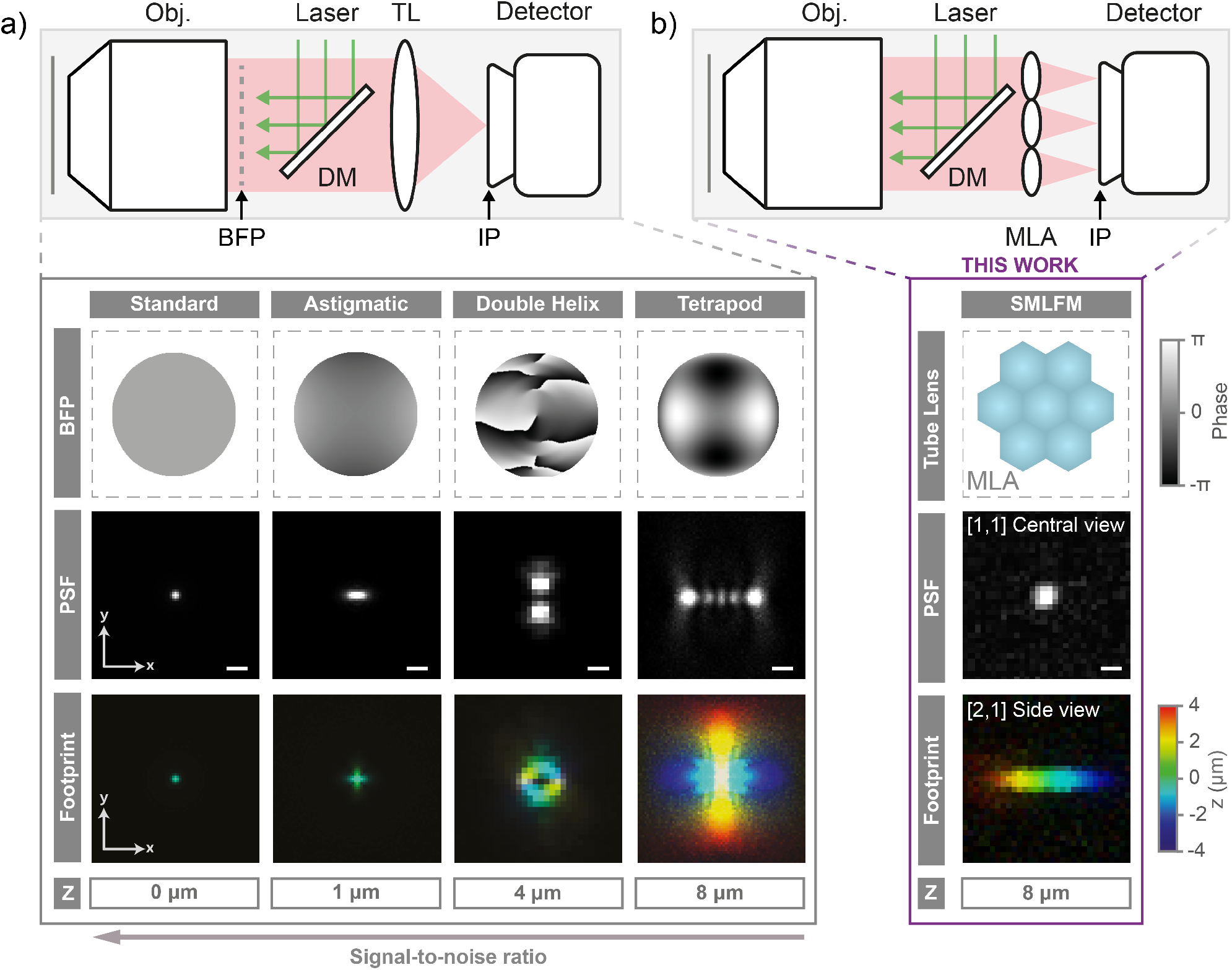
Encoding the 3D position of single molecule fluorescence into a 2D image. **a)** Optical schematic of a typical widefield microscope, where Obj. = objective lens, TL = tube lens, IP = image plane, BFP = back focal plane, and DM = dichroic mirror. Below are some common 3D point spread functions (PSFs) implemented by phase modulation in the BFP (top row), including the standard PSF, astigmatic PSF, double helix PSF and the tetrapod PSF (middle row) and their associated PSF footprints integrated over their entire axial range (bottom row). Field-of-view is 8 × 8 μm^−2^ and the scale bars are 1 μm. **b)** Optical schematic of a Fourier light field microscope for SMLFM, where MLA = microlens array (tube lens), which is placed in the BFP. Below is a schematic of the MLA, the PSF in the central perspective view, and the PSF footprint in the entire 8 μm axial range (see Supplementary Fig. S1 for further details). Pixel size is 110 nm for standard, astigmatic and the tetrapod PSF, and 266 nm for the DHPSF and SMLFM to reflect experimental parameters.

Raw localization datasets were simulated for the SMLM modalities presented in Fig. 1 (standard, astigmatic, double helix, light field and tetrapod) to investigate the effect the of PSF footprint on the ability to resolve single emitters at high densities (Fig. 2a). Briefly, the emitter density (ρ_loc_) of simulated SMLM data was systematically increased from 0.005 μm^−2^ (2 localizations per 20 μm × 20 μm field-of-view, FoV) to 0.375 μm^−2^ (150 localizations) and subsequently processed using conventional fitting algorithms (see Methods). Each dataset was also simulated for typical photon values expected for a fluorescent protein (1,000 photons), an organic dye (4,000 photons) and a next-generation fluorescent probe (10,000 photons) to reflect different labelling scenarios. Computational multi-emitter fitting was not implemented in the analysis to ensure a fair comparison across methods since algorithms are at different levels of technical development for each technique (21, 29), and single-emitter algorithms have been previously shown to outperform multi-emitter algorithms in high density 3D SMLM scenarios (12).

**Fig. 2.**
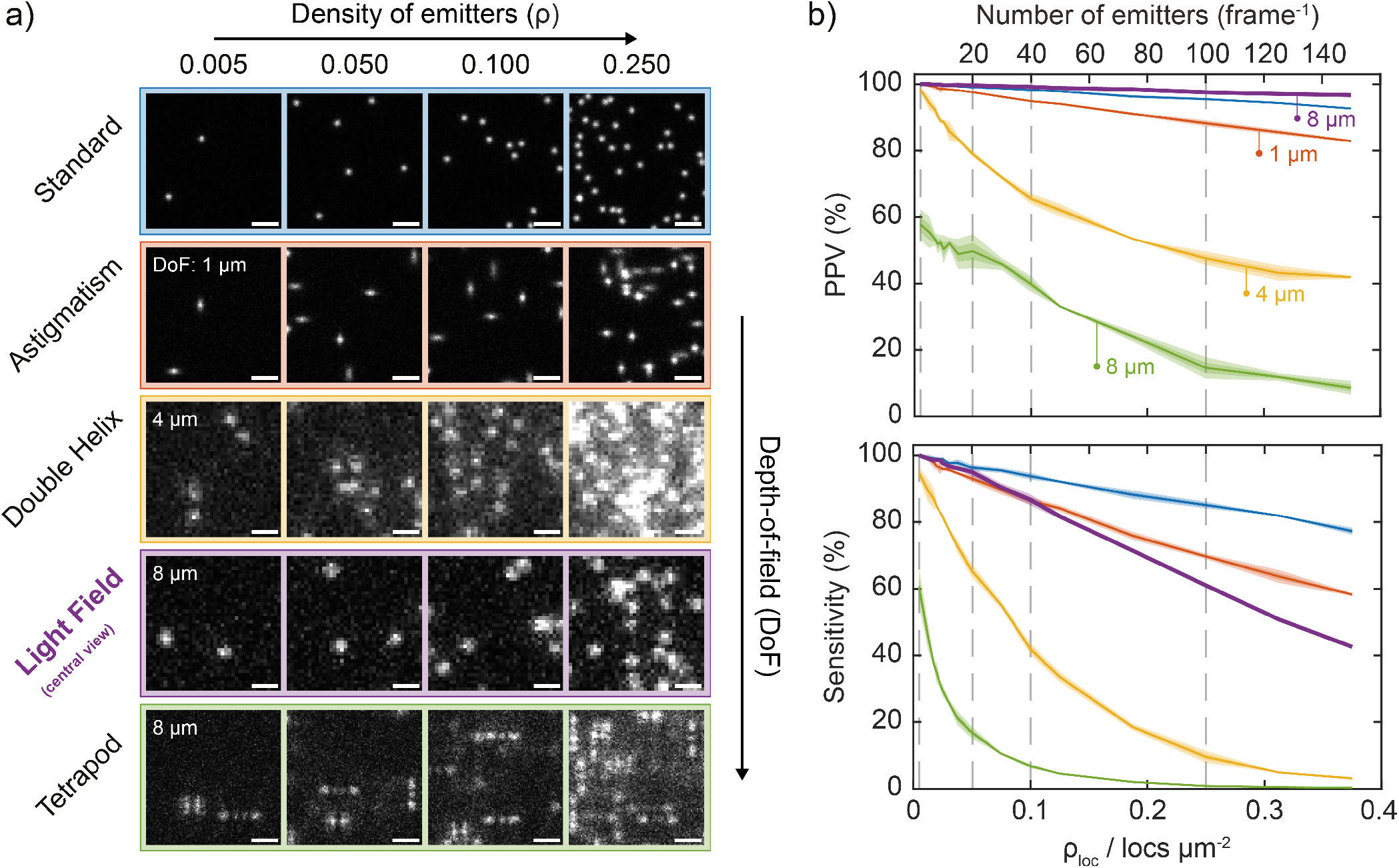
SMLFM consistently outperforms other 3D-SMLM techniques at correctly identifying and reconstructing single emitters at increasing densities. **a)** Snapshots of simulated raw localization datasets in a 10 × 10 μm^−2^ region for each imaging modality discussed herein (2D, astigmatism, double helix PSF, light field [central view] and tetrapod PSF.) Scale bar represents 2 μm. **b)** Top: Average positive predictive value (PPV) curves for each SMLM technique as a function of emitter density (ρ_loc_) at 4,000 detected photons, where PPV refers to the number of true positive localizations *vs*. total number of fitted localizations. Bottom: Average sensitivity curves as a function of ρ_loc_ at 4,000 detected photons, where sensitivity refers to the number of true positive localizations *vs*. total number of ground truth localizations. Shaded regions represent first and second standard deviation from the mean over three repeats of 100-frame simulated datasets. Example simulated datasets are presented in Supplementary Movie 1.

Direct comparison of PSF footprint with DoF (presented in Supplementary Fig. S3a) reveals how the area of each 3D PSF changes with axial position. The light field PSF is highly competitive, achieving an axial range suitable for imaging entire cells (8 μm) with a PSF area 55% the size of the DHPSF on average and 18% the area of the tetrapod PSF. As every perspective view comprises a super-resolvable image of the sample, only the area of the SMLFM PSF in each perspective view is needed for direct comparison. The simple and compact PSF footprint of SMLFM is a principle component in the ability to resolve single emitters at higher spot densities than the double helix and tetrapod PSFs. In this work we consider the central view for density studies.

Several quality-of-imaging metrics were then computed for each simulated dataset classifying a localization as either a true positive (TP), false positive (FP) or false negative (FN) with respect to known ground truth (GT) coordinates. The positive predictive value (PPV, also known as *precision*) describes the fraction of TP localizations relative to all localizations (TP+FP). Sensitivity (also known as *re-call*) describes the fraction of accurate localizations that are retrieved (TP/GT). Both PPV and sensitivity are presented as a function of ρ_loc_ in Fig. 2b using datasets simulated at 4,000 detected photons to reflect labelling using an organic dye molecule (see Supplementary Fig. S5 for PPV and sensitivity plots at 1,000, 4,000 and 10,000 detected photons).

Low signal-to-noise ratio (SNR), high background fluorescence and emitter overlap contribute to reconstruction artefacts from the incorrect localization of single emitters. This leads to a decrease in both PPV and sensitivity as a function of ρ_loc_, in agreement with similar work (6). Unlike for the double helix or tetrapod PSFs, the PPV for SMLFM is linear across the whole ρ_loc_ range with a maximum value of 100.0 ± 0.0% (mean ± SD) at ρ_loc_ = 0.005 μm^−2^ and a minimum of 96.7 ± 0.1% at ρ_loc_ = 0.375 μm^−2^, which can be rationalised by the spatio-temporally correlated PSF filtering out stochastic noise due to the requirement for the same emitter to be localized in each perspective view. PPV for the standard and astigmatic PSFs is also linear across all values of ρ_loc_ as expected from compact (but very low DoF) PSF footprints. Conversely, with an average pixel area of 1.8× that of the SMLFM PSF, the DHPSF exhibits a non-linear response to ρ_loc_ and a much lower PPV than SMLFM and likewise with the tetrapod PSF. Their weaker resistance to increasing ρ_loc_ can be attributed to their greater size and complexity of photon distributions, for example the tetrapod PSF was specifically designed for optimal Fisher information, and hence low density imaging scenarios (21, 30).

A linear relationship between sensitivity and ρ_loc_ is also observed for SMLFM with a maximum value of 100.0 ± 0.0% when ρ_loc_ = 0.005 μm^−2^ and a minimum of 42.5 ± 0.1% when ρ_loc_ = 0.375 μm^−2^. This is a result of distinguishing overlapping emitters through parallax, whereby single-molecule fluorescence is observed at different positions in each perspective view. Resolving overlapping emitters through the double helix and tetrapod PSF shaping methods is either impossible or computationally expensive during post processing. Alternatively, SMLFM facilitates these higher localization rates by resolving emitters through parallax, described herein as optical multi-emitter fitting (distinct from computational multi-emitter fitting). These data combined demonstrate that SMLFM has the capacity to localise 86.6 ± 0.9% of all emitters at a typical 2D-SMLM localization density of ∼0.1 μm^−2^ without compromising on the total number of localizations (31). Even at an incredibly high localization density of 0.375 μm^−2^ SMLFM is able to recover 43% of all ground truth localisations while this is less than 1% for the double helix and tetrapod PSFs.

By comparing the spot densities at which the DHPSF and SMLFM achieve equal error rates we determine a maximum speed improvement of 8.95× for SMLFM at an error rate of 13.5% (at which 86.5% of all localisations are correctly reconstructed in 3D, see Supplementary Note S4 section S4.2 and Supplementary Fig. S4) for 4,000 detected photons. This represents the upper practical limit in what SMLFM can achieve in direct comparison with the DHPSF (the state-of-the-art 3D SMLM modality for DoF and localization precision.) Furthermore, this maximum practical speed improvement was measured to be 25.3× (error rate of 40.0%) at 1,000 detected photons and 10.6× (error rate of 11.0%) at 10,000 photons. Therefore, on the basis of speed, SMLFM significantly out-performs the DHPSF at all light levels, particularly at low SNR, aided by the division of background photons across seven lenses.

### SMLFM captures the heterogeneity of live B-cell membrane receptors

The density-dependence studies reveal an optical redundancy in SMLFM that would be suited to the high-density volumetric imaging of entire cells through optical multi-emitter fitting. To challenge our method we imaged whole primary mouse B-cell membranes in a scan-free dSTORM modality previously optimised for 2D-SMLM (32–34). The 3D organisation of membrane receptors on immune cells, such as the B-cell receptor (BCR) is of increasing scientific interest to better understand the immune response to infection (35, 36). Single BCR complexes were labelled with a single molecule of Alexa-Fluor 647 (see Methods) and imaged under an inclined illumination angle to improve contrast (see Supplementary Note S1 section S1.3). An average of 40,000 3D localizations were accumulated per cell over an axial range of ∼8 μm (Fig. 3a-d). Membrane ruffles and microvilli could be observed, consistent with sub-diffraction resolution being obtained (37).

**Fig. 3.**
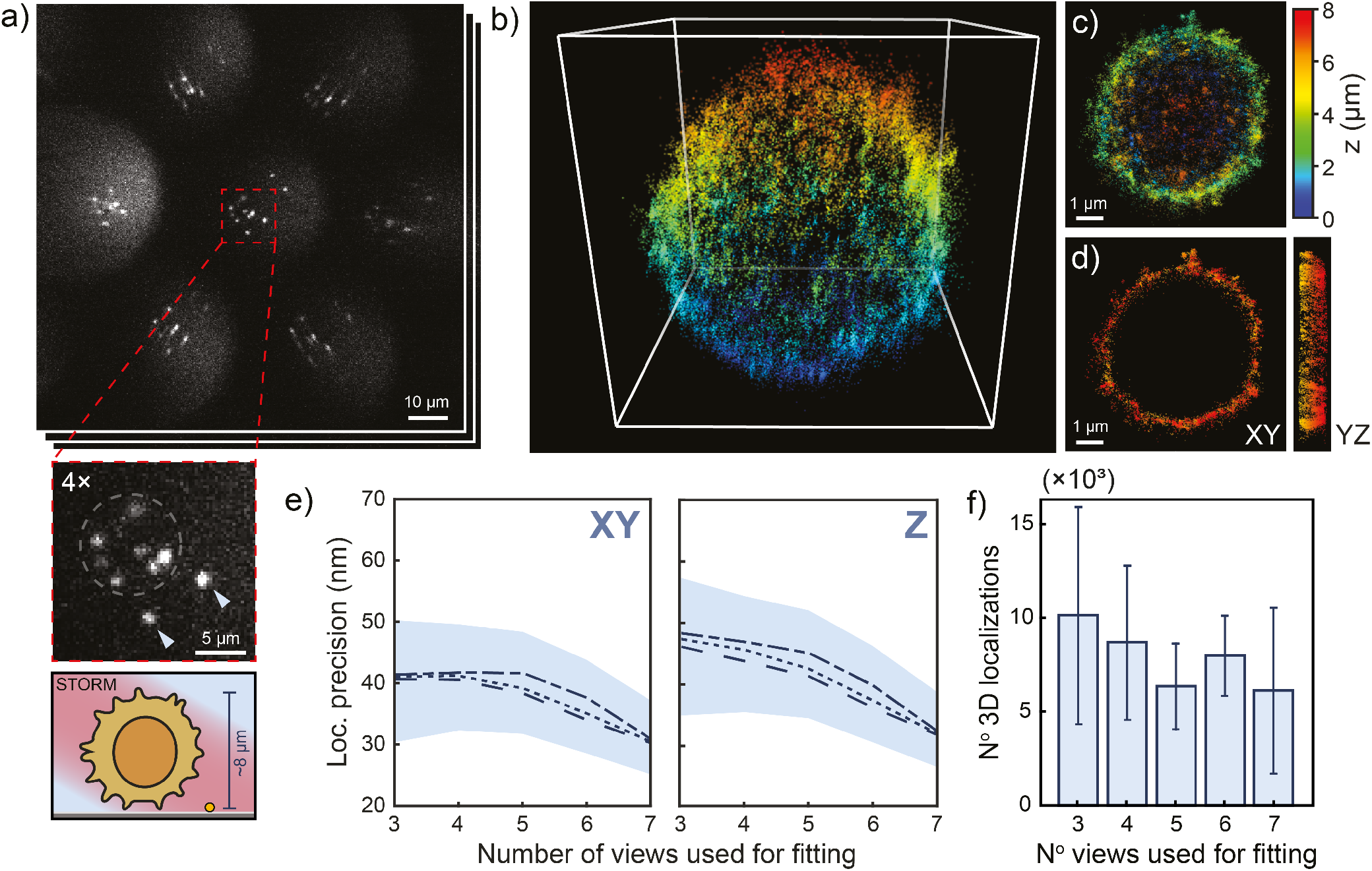
Scan-free SMLFM-STORM imaging of B cell receptors over whole primary mouse B cells. **a)** Raw SMLFM data of individual membrane receptors comprising 7 perspective views in a hexagonal arrangement. Expanded insert shows seven fluorescent puncta (Alexa Fluor 647) in the central perspective view and two fiducial markers indicated with arrows. Directly below is an illustration of the cell being imaged. **b)** Associated 3D reconstruction of the whole primary mouse B cell (40,000 3D localizations in a 9 μm^3^ box), **c)** an xy projection and **d)** a 1 μm^−2^ thick central clipping to illustrate non-internalisation of dye molecules. **e)** Median lateral and axial fitting error for localizations below 60 nm precision as a function of the number of views used to reconstruct a 3D localization (shading represents interquartile range). **f)** Associated proportion of 3D localizations below 60 nm lateral precision as a function of the number of views used to reconstruct a 3D coordinate.

3D localizations were collected over a ∼50 μm^2^ circular detector area (image space) with an average localization density of ∼0.10 μm^−2^ (see Supplementary Fig. S7) corresponding to a PPV of 99.2 ± 0.1% and sensitivity of 86.6 ± 0.9%. A maximum localization density of ∼0.24 μm^−2^ was achieved for a small portion of the experiment, which corresponds to a PPV of 97.6 ± 0.0% and sensitivity of 61.0 ± 0.2%. In comparison, at an average density of ∼0.10 μm^−2^ the DHPSF would be expected to achieve a PPV of 65.7 ± 1.4% and a sensitivity of 24.0 ± 0.5%, accurately localizing a quarter of all emitters. Equally, the tetrapod PSF would be expected to correctly localize 39.9 ± 1.2% of all emitters and recover 6.8 ± 0.4% of all localizations. The pronounced improvement in performance shown here by SMLFM at high localization densities, in addition to the imaging of complete cellular volumes without scanning, enables a significant improvement in sample throughput in future 3D-SMLM experiments.

SMLFM is advantageous at high densities because single emitters are not required to be isolated in every perspective view to be localized in 3D. We quantified this redundancy that enables optical multi-emitter fitting in these large cellular datasets by considering the localization precision as a function of number of perspective views used for PSF fitting. Fig. 3e reveals excellent localization precisions of ∼40 nm laterally (median) and ∼47 nm axially and these values improved to ∼30 nm laterally and ∼34 nm axially as the number of perspective views for fitting was systematically increased from 3 to 7. This is consistent with a higher effective numerical aperture and better sampling of the PSF position when utilizing a greater number of views. Attempts were made to ensure the distribution of localizations per number of views was equal, see Fig. 3f. Taken together, these data show that optical multi-emitter fitting via parallax is a powerful approach to 3D localizing single molecules at high densities within cells.

Another important application of the high emitter density measurements afforded by SMLFM is single-particle tracking (SPT) (38–40). 3D-SPT better quantifies diffusive processes than 2D measurements, which tend to underestimate diffusion rates (41, 42). A previous study of membrane protein mobility highlighted the importance of imaging diffusion dynamics away from the glass interface (basal surfaces) (39), which sparked the imaging of apical surfaces in 4 μm optical sections using the DHPSF (5). SMLFM boasts a significant practical advancement over this work, which is two-fold:

1. The larger DoF ensures single proteins can be tracked over entire cell volumes without scanning.
2. Localizing emitters through parallax improves the ability to delineate trajectories that would otherwise be occluded at higher densities.

To demonstrate this we applied SMLFM to the 3D-SPT of BCR complexes found on the surface of live mouse B cells, accumulating hundreds of trajectories in <10,000 frames (∼5 minutes) with an average track length of 12.5 points (Fig. 4a). Maximum likelihood estimation of the diffusion coefficient from trajectories over 5 cells (a total of 1806 tracks) yielded a distribution of diffusion coefficients (Fig. 4b-d) for individual BCR complexes with a median value of 0.14 ± 0.08 μm^2^ s^−1^, consistent with that observed by Tolar *et al*. on resting murine B cells (43). To confirm this, we measured a median diffusion coefficient of 0.20 ± 0.01 μm^2^ s^−1^ at the apical surface using fluorescence correlation spectroscopy.

**Fig. 4.**
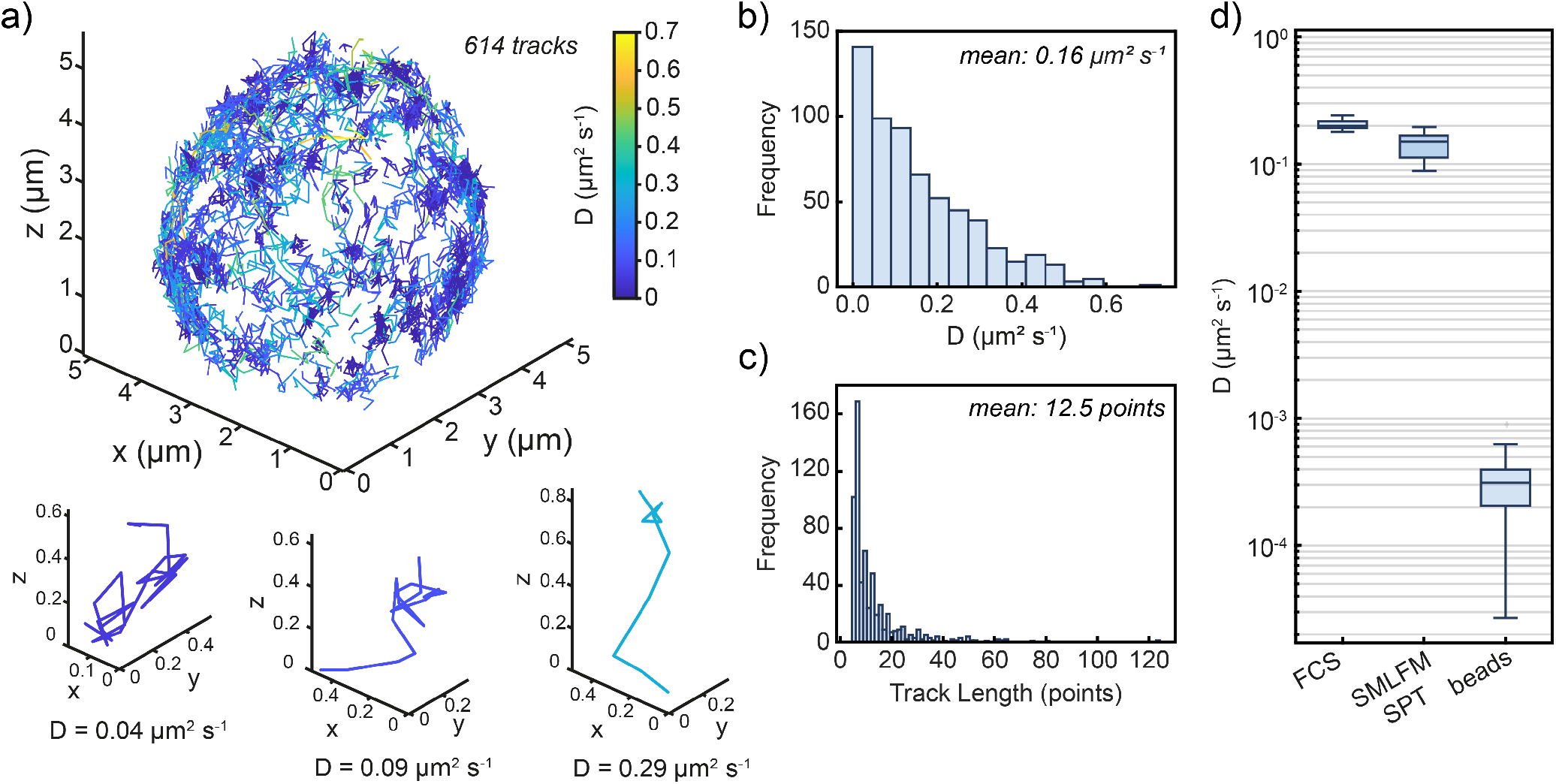
Scan-free whole-cell 3D SPT of the B cell receptor on primary mouse B-cell membranes using SMLFM. **a)** 3D trajectory map of the BCR over a whole primary mouse B cell totalling 614 tracks color-coded by diffusion coefficient using maximum likelihood estimation. Example isolated trajectories of varying diffusion coefficient are expanded directly below. Histogram of **b)** associated diffusion coefficients and **c)** track lengths (bin widths were determined using Freedman-Diaconis’ rule). **d)** Median diffusion coefficient of the BCR measured by FCS (N = 7) and SMLFM-SPT (N = 5), and the minimum diffusion coefficient measurable by SMLFM-SPT determined with immobilised beads (N = 9). SMLFM-SPT comprises a total of 1806 trajectories over 5 cells with a mean track length of 8 points.

SMLFM effectively and accurately captures the heterogeneity of diffusion coefficients of surface receptors and opens up the possibility of the direct observation of dynamic BCR clustering following or proceeding antigen encounters (and in general the clustering of key signalling proteins in other systems), which is of great interest in the study of receptor triggering (44).

### High-density SMLFM resolves intracellular structure

To demonstrate intracellular imaging at very high emitter density, we performed scan-free dSTORM imaging of tubulin in fixed HeLa cells with SMLFM. Fig. 5a contains a snapshot of raw localization data with a cell occupying a 40 μm × 40 μm FoV, which spanned an axial range of ≤ 3μm, with the 3D reconstruction shown in Fig. 5b. An expanded region is presented in Fig. 5c. Line profiles (Figure 5d, width 400 nm) were drawn for two ranges to confirm the resolution of individual microtubules. A maximum of 40 3D localizations were detected per image frame, with an average of ∼22 per frame, totalling 150,000 localizations over ∼4 minutes (30 ms detector exposure, Fig. 5e). This corresponds to a maximum density of 0.15 μm^−2^ and an average of 0.075 μm^−2^, whereupon through simulations SMLFM is shown to retrieve 90.4 ± 0.6% of all localizations (sensitivity) with an accuracy of 99.4 ± 0.0% (PPV). A median of 3,900 photons were detected per 3D localization (Fig. 5f) achieving median lateral and axial localization precisions of 42.6 nm and 46.7 nm, respectively (Supplementary Fig. S10). A Fourier Shell Correlation (FSC) of 54 nm resolution at a 1/7 cut-off was calculated from the localizations presented in Figure 5c.

**Fig. 5.**
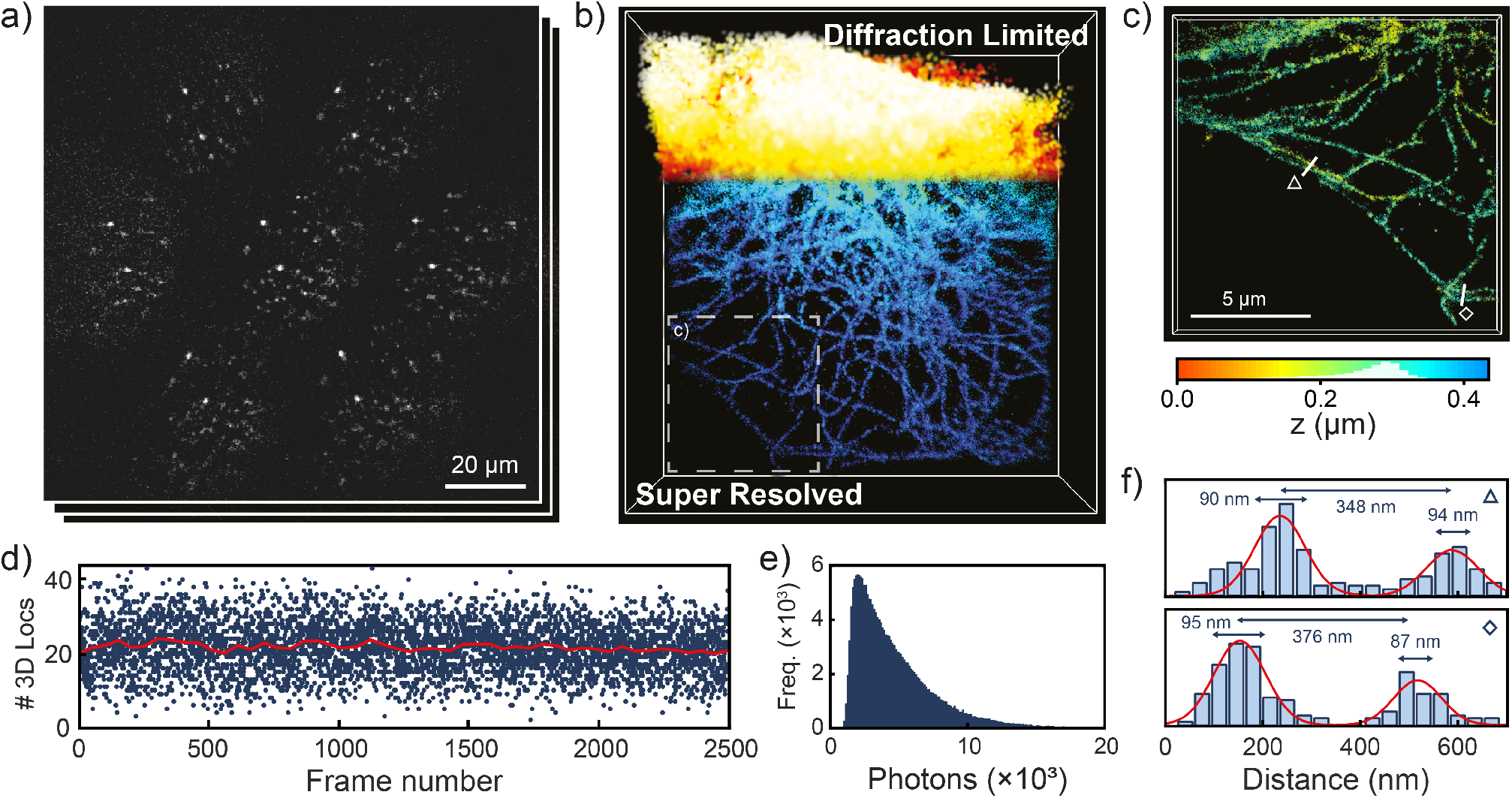
SMLFM-STORM imaging of Alexa Fluor 647-labelled tubulin in a HeLa cell. **a)** A snapshot of raw localization data in microtubule-stained HeLa cells imaged through the 7-hex SMLFM platform and **b)** the corresponding super-resolved 3D volume. The super-resolved area is 23 μm× 12 μm in size and contains 150,000 3D localizations. **c)** Expansion of the marked region (60,000 3D localizations) in **b). d)** localization rate over the first 2500 frames indicating a mean 3D localization rate (red line, rolling average over 100 frames) of 22 frame^−1^ and an upper limit of ∼40 frame^-1^ corresponding to ∼0.075 and ∼0.15 locs μm^-2^ respectively. **e)** Histogram of detected photons per 3D localization (a median value of 3,900). **f)** Line plots through pairs of microtubules (width of 400 nm) in panel **d)** showing individual microtubules being resolved.

Furthermore, no 3D-specific sample optimisation was undertaken prior to imaging and a dSTORM buffer protocol was implemented that was previously developed for 2D-SMLM (33). As protocol optimisation for SMLM can be time intensive and challenging, this facile translation from 2D to 3D SMLM presents a significant advantage in the future use of 3D-SMLM in biological research (45).

## Conclusions

We report the first hexagonal SMLFM platform enabling 3D-SMLM over an 8 μm axial range and quantitatively compared its performance to other common 3D PSFs through simulations revealing an order-of-magnitude speed improvement compared to DHPSF microscopy. We attribute this speed improvement to optical multi-emitter fitting through which overlapping emitters in an imaging volume can now be resolved through a redundancy in the number of perspective views required for 3D reconstruction. We applied SMLFM experimentally to the imaging in both live and fixed whole cells and dense arrays of cytosolic tubulin, where it consistently localised single emitters in 3D at high, non-optimised, densities achieving localization precisions ∼40 nm laterally and ∼50 nm axially.

Future endeavours could couple SMLFM with computational multi-emitter fitting and/or deep learning strategies, high speed detectors, and alternative volumetric labelling strategies to push localization rates even further. We anticipate the uptake of SMLFM as a powerful tool in improving our understanding of 3D nano-scale architecture, dynamics and will bring robust real-time 3D super-resolution imaging to life scientists.

## Methods

### Optical setup

The SMLFM platform described in this work was constructed using an epi-fluorescence microscope (Eclipse Ti-U, Nikon) housing a 1.27 NA water immersion objective lens (Plan Apo VC 60×, Nikon, Tokyo, Japan) for imaging above the coverslip. The z-position of the objective was controlled with a scanning piezo (P-726 PIFOC, PI, Karlsruhe, Germany). The Fourier lens (f = 175 mm, ThorLabs) was placed in a 4*f* configuration with the tube lens (f = 200 mm, Nikon) to relay the back focal plane (BFP) outside of the microscope body (see Supplementary Fig. S1). A hexagonal microlens array (f = 175 mm, pitch = 2.39 mm, custommade by CAIRN) was placed in the BFP to relay the image plane onto an EMCCD (Evolve Delta 512, Photometrics, Tucson, AZ, 16 μm pixel size). Excitation was achieved using a 640 nm (∼10 kW cm^−2^ power density, 150 mW, iBeam Smart-S 640-S, Toptica, Munich, Germany) and activation by a 405 nm (∼0.04 kW cm^−2^ power density, 120 mW, iBeam Smart-S 405-S, Toptica, Munich, Germany) laser, that were circularly polarised, collimated and focused on to the BFP of the objective to create an evanescent excitation wave. Unless stated otherwise, samples were excited with a highly inclined and laminated optical sheet (HILO) which was achieved by laterally displacing the excitation beam towards the edge of the BFP of the objective (see Supplementary Note S1 section S1.3). Fluorescence was collected by the same objective and separated from the excitation beam using a quad-band dichroic mirror (Di01-R405/488/561/635-25×36, Semrock, Rochester, NY). Long-pass (BLP02-640R-25, Semrock) and bandpass (FF01-680/X-25, Semrock) emission filters were placed immediately before the detector to isolate fluorescence emission. The pixel size in image space was measured at 266.

### 3D reconstruction of SMLFM data

All experimental data were recorded as .*tif* stacks. 2D gaussian fitting of all emitter positions in all perspective views was carried out in Fiji using PeakFit (GDSC SMLM 2.0) to yield a set of 2D localizations for each raw frame. Given this initial set of 2D localizations, individual emitters were localized in 3D using custom Matlab scripts as outlined in (22). Briefly, the most likely subset of 2D localizations in different perspective views corresponding to a unique emitter were identified. Provided that this set of localizations contained more than 3 elements, the 3D location of this emitter was calculated as the least-squares estimate to an optical model relating axial emitter position to the parallax between perspective views. If the residual light field fit error was below 200 nm, the fit was accepted and the subset of 2D localizations was removed. This procedure was repeated for each individual emitter. Drift correction was performed by localizing the position of a fiducial marker in each frame and subtracting the resulting 3D fiducial points from all localizations of the corresponding frame. System and sample aberrations were corrected for by subtracting the residual disparity (calculated for data acquired for all emitters localized during the first 1,000 frames) from all 2D localizations prior to calculating the 3D light field fit. For full details of the light field localization fitting procedure refer to the Supplementary Information of (22). 3D visualisation was carried out in ViSP (46).

### Optical 3D calibration

Fluorescent beads (200 nm, Deep Red FluoSpheres, ThermoFisher, Waltham, MA) were immobilised on a glass slide and imaged to calibrate for deviations in experimental and calculated the disparity from the SMLFM optical model. Glass slides were cleaned under argon plasma (PDC-002, Harrick Plasma, Ithaca, NY) for 1 hour and incubated with poly-L-lysine (PLL, 50 μL, 0.1% w/v, Sigma-Aldrich, P820) for 10 minutes. Glass slides were washed with PBS (3 × 50 μL) and incubated with fluorescent beads (50 μL, ca. 3.6 × 108 particles/mL) incubated for 3 minutes before washing further with PBS (3 × 50 μL). The piezo stage (P-726 PIFOC, PI, Karlsruhe, Germany) was used to scan the objective lens axially over 8 μm recording 10 frames at 30 ms exposure per 60 nm increment. The data was reconstructed in 3D and plotted against the known movement of the piezo stage. A linear fit was applied to the calibration curve, the gradient of which was a correction factor subsequently applied to all reconstructed data presented in this work.

### SPT analysis

Following 3D reconstruction of SMLFM data a custom-written MATLAB code was implemented to temporally group localizations into single trajectories. Some parameters were chosen by the user, including number of dark frames, linking distance, and minimum track length. The diffusion coefficient was then calculated from each trajectory using maximum likelihood estimation, which has previously been shown to yield statistically robust measurements of the diffusion coefficient (47).

To determine the minimum observable diffusion coefficient, fluorescent beads (200 nm, Deep Red FluoSpheres, ThermoFisher, Waltham, MA) were immobilised on a glass slide and imaged under conditions (641 nm excitation at ∼2 mW cm−2 power density, 20 ms exposure time) that artificially reproduce the same photon intensities as PA-JF646 used for SPT experiments. The raw data was reconstructed in 3D and trajectories analysed as described previously (22) to yield the smallest resolvable diffusion coefficient.

### Analysis of simulated data

2D and astigmatic datasets were fitted in PeakFit (GDSC SMLM 2.0, Fiji plug-in) using a circular and astigmatic Gaussian PSF, respectively. DHPSF datasets were initially fitted in PeakFit using a circular Gaussian PSF before 3D reconstruction using DHPSFU (https://github.com/TheLaueLab/DHPSFU). SMLFM datasets were initially fitted in PeakFit using a circular Gaussian PSF before 3D reconstruction using a custom MATLAB code described previously. Tetrapod PSF data was fitted using Zola (Fiji plug-in) for 3D reconstruction (48).

A custom MATLAB code was written to compare the fitted (3D, 2D for the standard PSF) point data to the ground truth coordinates. Specifically, the root mean square distance matrix is calculated between all ground truth coordinates and all reconstructed data points on a frame-by-frame basis and counted as either a true positive, false positive or false negative given a user-specified distance tolerance. The tolerance applied was different for each technique and dictated by the precision and thresholds (determined by the fitting error) were applied to determine true positive and false positives.

### Preparation of coverslips for B-cell imaging

Glass slides (24 × 50 mm borosilicate, thickness No. 1, Brand, Wertheim, Germany) were washed with propan-2-ol and water, dried under nitrogen and cleaned under argon plasma (PDC-002, Harrick Plasma, Ithaca, NY) for 1 hour. Glass slides were then incubated with poly-L-lysine (PLL, 50 μL, 0.1% w/v, Sigma-Aldrich, P820) for 1 hour and washed with filtered (0.02 μm syringe filter, Whatman, 6809-1102) PBS (3 × 50 μL) before incubation with gold nanoparticles (5 μL, 0.1 μm, Merck) for 20 minutes.

For fixed cell imaging, glass slides were then washed with filtered PBS (3 × 50μL). 1 × 105 fixed labelled B cells were washed in dSTORM buffer (50 mM Tris-HCl, 10 mM NaCl, 10% glucose, 10 mM MEA, 84 μg/mL catalase, 0.2 mg/mL GLOX, adjusted to pH 8), plated in 20 μL dSTORM buffer and left to settle for *>* 20 minutes. Prior to imaging, the sample was washed into fresh buffer dSTORM buffer.

For live cell tracking, PLL-coated glass slides were prepared as above and placed in filtered PBS. For SPT, the surface was incubated with gold beads as above, washed 3× in filtered PBS, and cells labelled with PA-JF646-conjugated Fab-Halo were allowed to settle onto the surface for 5–10 minutes prior to imaging.

For point fluorescence correlation microscopy (pFCS), cells labelled with AF647-conjugated Fab-HaloTag were incubated onto the PLL surface for 5–10 minutes and imaged using an Zeiss LSM780 inverted confocal microscope using a 40× water objective, with the sample excited using a 633 nm He-Ne laser. The confocal volume was placed on the apical surface of the cell membrane and five repeated measurements were taken per cell. Data was analysed using PyCorrFit and the diffusion coefficient calculated from the average transit time (*τ*), using the confocal beam width as calculated using a solution of 100 nM AF647 HaloTag ligand solution.

### B-cell culture and fluorescent labelling

Primary murine B cells were isolated from the spleens of male C57BL/6J mice aged between 8 and 12 weeks. Splenocytes were isolated by mechanical disruption of the spleen, and incubated with ACK lysing buffer (Lonza, LZ10-548E) for 2 minutes at room temperature to lyse erythrocytes. The cells were washed in RPMI-1640 (Gibco) medium supplemented with 10% foetal bovine serum (FBS) and B cells were isolated using a B Cell Isolation Kit, mouse (Miltenyi Biotec, 130-090-862) according to the manufacturer’s instructions. Purified murine B cells were either resuspended in PBS for dSTORM labelling or frozen in FBS supplemented with 10% DMSO to later culture for live cell imaging (SPT and FCS).

BCR complexes were labelled using a recombinant protein based on the Fab fragment of the anti-murine CD79b antibody HM79-16. A self-labelling HaloTag domain was introduced to the C-terminus of the Fab heavy chain to ensure single-dye labelling of the probe. Fab-Halo protein was labelled with HaloTag ligand dyes by incubation with 2-fold molar excess of dye for 90 minutes at room temperature, with free dye removed using a Bio-Spin P-6 gel column (BioRad, 7326227) according to manufacturer’s instructions. Labelled protein was aliquoted and stored at −80 °C. For dSTORM imaging, freshly isolated C57BL/6J B cells were labelled at 4 °C with recombinant Alexa Fluor 647 Fab-Halo protein. 2 × 106 cells were washed in 0.22 μm−2 filtered PBS and incubated in 2.5 μM Fab-Halo (AF647) for 45 minutes at 4 °C. Cells were washed twice in cold filtered PBS, fixed in 1% paraformaldehyde (Sigma, 28906) for 30 minutes at 4 °C, and placed in filtered PBS at a final density of 4 × 107 cells/mL.

For live cell imaging, as conducted for SPT and FCS, cells were thawed from frozen stocks and cultured in primary B cell medium (RPMI-1640 supplemented with 10% FBS, 2mM L-Glutamine, 10 mM HEPES, 1 mM sodium pyruvate, 50 μM 2-mercaptoethanol, 50 U/mL penicillin and 50 μg/mL streptomycin), supplemented with 10 μg/mL anti-mouse CD40 (clone 1C10, Biolegend 102812) and 10 ng/mL murine IL-4 (Peprotech, 214-14). For live cell imaging, 2 × 105 cells were washed in filtered PBS and incubated with 1 μM fluorescent Fab-Halo for 15 minutes at room temperature, and washed twice in PBS prior to incubation with the coverslip.

### HeLa cell culture and fluorescent labelling

HeLa TDS cells were cultured at 37 °C and 5% CO2 in DMEM (Gibco, Invitrogen) supplemented with 10% FBS (Life Technologies), 1% penicillin/streptomycin (Life Technologies), and 1% glutamine (Life Technologies). Cells were passaged every three days and were regularly tested for mycoplasma. One day prior to fixation, cells were seeded on high-precision 1.5 glass coverslips (MatTek, P35G-0.170-14-C) for imaging.

Cells were fixed and permeabilised simultaneously for 6 minutes in Cytoskeleton Buffer with Sucrose (CBS, 10 mM MES, 138 mM KCl, 3 mM MgCl2, 2 mM EGTA, and 4.5% sucrose w/v, pH 7.4) containing 4% methanol-free formaldehyde (FA, Fisher Scientific) and 0.2% Triton, followed by a second fixation for 14 minutes in CBS + 4% methanol-free formaldehyde at 37 °C and 5% CO2. Post fixation, cells were washed three times in PBS + 0.1% Tween (PBST), and further permeabilised in PBS + 0.5% Triton for 5 minutes. Cells were then washed in PBST three times and blocked in 5% BSA (in PBS) for 1 hour at RT. Samples were further washed three times in PBST, after which samples were incubated with an anti-*α*-tubulin antibody (ab7291, clone DM1A, at 2.5 μg mL−1 in 5% non-fat milk) overnight at 4 °C. Cells were then washed six times in PBST after which a Donkey anti-Mouse IgG (H+L) Highly Cross-Adsorbed Secondary Antibody AlexaFluor 647 (Invitrogen, A-31571, at 2.0 μg mL−1 in 5% non-fat milk) was added to the sample for 1 hour at 4 °C. The cells were then washed six times in PBS and the sample flooded with STORM imaging buffer prepared as described previously.

## ACKNOWLEDGEMENTS

The authors would like to thank Jeremy Graham at CAIRN Research for providing the hexagonal microlens array. The authors thank Alexander Collins for useful discussions and Gregory Chant and James McColl for assisting with the optimisation of the dSTORM buffer. We thank Janelia Materials for providing PA-JF646 Halo-Tag used for 3D-SPT.

## AUTHOR CONTRIBUTIONS

SD, AH and SFL conceived the project. SFL supervised the research. SD and AH built the optical set-up and performed baseline experiments. SD performed all SMLFM imaging and data analysis. JFF prepared all labelled B-cell samples and conducted FCS experiments. EB programmed and performed the simulations. RP and BZ prepared labelled tubulin samples. ES provided samples for early tests. RRS, KOH, SB and BZ wrote and maintained the 3D reconstruction code. JSB provided diffusion analysis code. SD and SFL wrote the manuscript with input from all authors.

## References

1. Rust, M.J., Bates, M. and Zhuang, X. Sub-diffraction-limit imaging by stochastic optical reconstruction microscopy (STORM). Nature Methods, 3(10):793–796, October 2006. ISSN 1548-7105. doi:10.1038/nmeth929. Number: 10 Publisher: Nature Publishing Group.

2. Betzig, E., Patterson, G.H., Sougrat, R., Lindwasser, O.W., Olenych, S., Bonifacino, J.S., Davidson, M.W., Lippincott-Schwartz, J. and Hess, H.F. Imaging Intracellular Fluorescent Proteins at Nanometer Resolution. Science, 313(5793):1642–1645, September 2006. ISSN 0036-8075, 1095-9203. doi:10.1126/science.1127344. Publisher: American Association for the Advancement of Science Section: Report.

3. Hess, S.T., Girirajan, T.P. and Mason, M.D. Ultra-high resolution imaging by fluorescence photoactivation localization microscopy. Biophysical Journal, 91(11):4258–4272, December 2006. doi:10.1529/biophysj.106.091116.

4. Lelek, M., Gyparaki, M.T., Beliu, G., Schueder, F., Griffié, J., Manley, S., Jungmann, R., Sauer, M., Lakadamyali, M. and Zimmer, C. Single-molecule localization microscopy. Nature Reviews Methods Primers, 1(1):39, December 2021. ISSN 2662-8449. doi:10.1038/s43586-021-00038-x.

5. Carr, A.R., Ponjavic, A., Basu, S., McColl, J., Santos, A.M., Davis, S., Laue, E.D., Klenerman, D. and Lee, S.F. Three-Dimensional Super-Resolution in Eukaryotic Cells Using the Double-Helix Point Spread Function. Biophysical Journal, 112(7):1444–1454, April 2017. ISSN 00063495. doi:10.1016/j.bpj.2017.02.023.

6. Nehme, E., Freedman, D., Gordon, R., Ferdman, B., Weiss, L.E., Alalouf, O., Naor, T., Orange, R., Michaeli, T. and Shechtman, Y. DeepSTORM3d: dense 3d localization microscopy and PSF design by deep learning. Nature Methods, 17(7):734–740, June 2020. doi:10.1038/s41592-020-0853-5.

7. Chen, B.C., Legant, W.R., Wang, K., Shao, L., Milkie, D.E., Davidson, M.W., Janetopoulos, C., Wu, X.S., Hammer, J.A., Liu, Z., English, B.P., Mimori-Kiyosue, Y., Romero, D.P., Ritter, A.T., Lippincott-Schwartz, J., Fritz-Laylin, L., Mullins, R.D., Mitchell, D.M., Bembenek, J.N., Reymann, A.C. et al. Lattice light-sheet microscopy: Imaging molecules to embryos at high spatiotemporal resolution. Science, 346(6208), October 2014. doi:10.1126/science.1257998.

8. Legant, W.R., Shao, L., Grimm, J.B., Brown, T.A., Milkie, D.E., Avants, B.B., Lavis, L.D. and Betzig, E. High-density three-dimensional localization microscopy across large volumes. Nature Methods, 13(4):359–365, March 2016. doi:10.1038/nmeth.3797.

9. Holden, S.J., Uphoff, S. and Kapanidis, A.N. DAOSTORM: an algorithm for highdensity super-resolution microscopy. Nature Methods, 8(4):279–280, April 2011. ISSN 1548-7091,1548-7105. doi:10.1038/nmeth0411-279.

10. Gu, L., Sheng, Y., Chen, Y., Chang, H., Zhang, Y., Lv, P., Ji, W. and Xu, T. High-Density 3D Single Molecular Analysis Based on Compressed Sensing. Biophysical Journal, 106 (11):2443–2449, June 2014. ISSN 0006-3495. doi:10.1016/j.bpj.2014.04.021.

11. Speiser, A., Müller, L.R., Hoess, P., Matti, U., Obara, C.J., Legant, W.R., Kreshuk, A., Macke, J.H., Ries, J. and Turaga, S.C. Deep learning enables fast and dense singlemolecule localization with high accuracy. Nature Methods, 18(9):1082–1090, September 2021. ISSN 1548-7105. doi:10.1038/s41592-021-01236-x. Number: 9 Publisher: Nature Publishing Group.

12. Sage, D., Pham, T.A., Babcock, H., Lukes, T., Pengo, T., Chao, J., Velmurugan, R., Herbert, A., Agrawal, A., Colabrese, S., Wheeler, A., Archetti, A., Rieger, B., Ober, R., Hagen, G.M., Sibarita, J.B., Ries, J., Henriques, R., Unser, M. and Holden, S. Superresolution fight club: assessment of 2D and 3D single-molecule localization microscopy software. Nature Methods, 16(5):387–395, May 2019. ISSN 1548-7091, 1548-7105. doi:10.1038/s41592-019-0364-4.

13. Grover, G., Quirin, S., Fiedler, C. and Piestun, R. Photon efficient double-helix PSF microscopy with application to 3D photo-activation localization imaging. Biomedical Optics Express, 2(11):3010–3020, November 2011. ISSN 2156-7085. doi:10.1364/BOE.2.003010. Publisher: Optical Society of America.

14. Rehman, S.A., Carr, A.R., Lenz, M.O., Lee, S.F. and O’Holleran, K. Maximizing the field of view and accuracy in 3D Single Molecule Localization Microscopy. Optics Express, 26 (4):4631–4637, February 2018. ISSN 1094-4087. doi:10.1364/OE.26.004631. Publisher: Optical Society of America.

15. Huang, B., Wang, W., Bates, M. and Zhuang, X. Three-dimensional Super-resolution Imaging by Stochastic Optical Reconstruction Microscopy. Science (New York, N.Y.), 319 (5864):810–813, February 2008. ISSN 0036-8075. doi:10.1126/science.1153529.

16. Backer, A.S., Backlund, M.P., von Diezmann, A.R., Sahl, S.J. and Moerner, W.E. A bisected pupil for studying single-molecule orientational dynamics and its application to three-dimensional super-resolution microscopy. Applied Physics Letters, 104(19):193701, May 2014. ISSN 0003-6951. doi:10.1063/1.4876440. Publisher: American Institute of Physics.

17. Lew, M.D., Lee, S.F., Badieirostami, M. and Moerner, W.E. Corkscrew point spread function for far-field three-dimensional nanoscale localization of pointlike objects. Optics Letters, 36(2):202–204, January 2011. doi:10.1364/OL.36.000202. Publisher: Optica Publishing Group.

18. Pavani, S.R.P., Thompson, M.A., Biteen, J.S., Lord, S.J., Liu, N., Twieg, R.J., Piestun, R. and Moerner, W.E. Three-dimensional, single-molecule fluorescence imaging beyond the diffraction limit by using a double-helix point spread function. Proceedings of the National Academy of Sciences, 106(9):2995–2999, March 2009. ISSN 0027-8424, 1091-6490. doi:10.1073/pnas.0900245106. Publisher: National Academy of Sciences Section: Physical Sciences.

19. Pavani, S.R.P. and Piestun, R. Three dimensional tracking of fluorescent microparticles using a photon-limited double-helix response system. Optics Express, 16(26):22048, December 2008. ISSN 1094-4087. doi:10.1364/OE.16.022048.

20. Shechtman, Y., Sahl, S.J., Backer, A.S. and Moerner, W. Optimal point spread function design for 3d imaging. Physical Review Letters, 113(13), September 2014. doi:10.1103/physrevlett.113.133902.

21. Shechtman, Y., Weiss, L.E., Backer, A.S., Sahl, S.J. and Moerner, W.E. Precise Three-Dimensional Scan-Free Multiple-Particle Tracking over Large Axial Ranges with Tetrapod Point Spread Functions. Nano Letters, 15(6):4194–4199, June 2015. ISSN 1530-6984. doi:10.1021/acs.nanolett.5b01396. Publisher: American Chemical Society.

22. Sims, R.R., Abdul Rehman, S., Lenz, M.O., Benaissa, S.I., Bruggeman, E., Clark, A., Sanders, E.W., Ponjavic, A., Muresan, L., Lee, S.F. and O’Holleran, K. Single molecule light field microscopy. Optica, 7(9):1065, September 2020. ISSN 2334-2536. doi:10.1364/OPTICA.397172.

23. Galdón, L., Saavedra, G., Garcia-Sucerquia, J., Martínez-Corral, M. and Sánchez-Ortiga, E. Fourier lightfield microscopy: a practical design guide. Applied Optics, 61(10):2558, March 2022. doi:10.1364/ao.453723.

24. Guo, C., Guo, C., Liu, W., Liu, W., Hua, X., Li, H. and Jia, S. Fourier light-field microscopy. Optics Express, 27(18):25573–25594, September 2019. ISSN 1094-4087. doi:10.1364/OE.27.025573. Publisher: Optical Society of America.

25. Abrahamsson, S., Chen, J., Hajj, B., Stallinga, S., Katsov, A.Y., Wisniewski, J., Mizuguchi, G., Soule, P., Mueller, F., Darzacq, C.D., Darzacq, X., Wu, C., Bargmann, C.I., Agard, D.A., Dahan, M. and Gustafsson, M.G.L. Fast multicolor 3D imaging using aberration-corrected multifocus microscopy. Nature Methods, 10(1):60–63, January 2013. ISSN 1548-7105. doi:10.1038/nmeth.2277. Number: 1 Publisher: Nature Publishing Group.

26. Prabhat, P., Ram, S., Ward, E. and Ober, R. Simultaneous imaging of different focal planes in fluorescence microscopy for the study of cellular dynamics in three dimensions. IEEE Transactions on NanoBioscience, 3(4):237–242, December 2004. ISSN 1558-2639. doi:10.1109/TNB.2004.837899. Conference Name: IEEE Transactions on NanoBioscience.

27. Ram, S., Prabhat, P., Ward, E.S. and Ober, R.J. Improved single particle localization accuracy with dual objective multifocal plane microscopy. Optics Express, 17(8):6881–6898, April 2009. ISSN 1094-4087. doi:10.1364/OE.17.006881. Publisher: Optica Publishing Group.

28. Juette, M.F., Gould, T.J., Lessard, M.D., Mlodzianoski, M.J., Nagpure, B.S., Bennett, B.T., Hess, S.T. and Bewersdorf, J. Three-dimensional sub–100 nm resolution fluorescence microscopy of thick samples. Nature Methods, 5(6):527–529, June 2008. ISSN 1548-7105. doi:10.1038/nmeth.1211. Number: 6 Publisher: Nature Publishing Group.

29. Backlund, M.P., Lew, M.D., Backer, A.S., Sahl, S.J., Grover, G., Agrawal, A., Piestun, R. and Moerner, W.E. The double-helix point spread function enables precise and accurate measurement of 3d single-molecule localization and orientation. In Enderlein, J., Gregor, I., Gryczynski, Z.K., Erdmann, R.and Koberling, F., editors, SPIE Proceedings. SPIE, February 2013. doi:10.1117/12.2001671.

30. Shechtman, Y., Sahl, S.J., Backer, A.S. and Moerner, W. Optimal Point Spread Function Design for 3D Imaging. Physical Review Letters, 113(13):133902, September 2014. ISSN 0031-9007, 1079-7114. doi:10.1103/PhysRevLett.113.133902.

31. Linde, S.v.d., Krstic, I., Prisner, T., Doose, S., Heilemann, M. and Sauer, M. Photoinduced formation of reversible dye radicals and their impact on super-resolution imaging. Photochemical & Photobiological Sciences, 10(4):499–506, March 2011. ISSN 1474-9092. doi:10.1039/C0PP00317D. Publisher: The Royal Society of Chemistry.

32. Sanders, E., Carr, A.R., Bruggeman, E., Koerbel, M., Benaissa, S.I., Donat, R., Santos, A.M., McColl, J., O’Holleran, K., Klenerman, D., Davis, S.J., Lee, S.F. and Ponjavic, A. resPAINT: Accelerating volumetric super-resolution localisation microscopy by active control of probe emission. Technical report, bioRxiv, April 2022. Section: New Results Type: article.

33. Lehmann, M., Lichtner, G., Klenz, H. and Schmoranzer, J. Novel organic dyes for multicolor localization-based super-resolution microscopy. Journal of Biophotonics, 9(1-2):161–170, 2016. ISSN 1864-0648. doi:10.1002/jbio.201500119. _eprint: https://onlinelibrary.wiley.com/doi/pdf/10.1002/jbio.201500119.

34. Gomes de Castro, M.A., Wildhagen, H., Sograte-Idrissi, S., Hitzing, C., Binder, M., Trepel, M., Engels, N. and Opazo, F. Differential organization of tonic and chronic B cell antigen receptors in the plasma membrane. Nature Communications, 10(1):820, February 2019. ISSN 2041-1723. doi:10.1038/s41467-019-08677-1. Number: 1 Publisher: Nature Publishing Group.

35. Treanor, B. B-cell receptor: from resting state to activate. Immunology, 136(1):21–27, May 2012. ISSN 1365-2567 0019-2805. doi:10.1111/j.1365-2567.2012.03564.x.

36. Rickert, R.C. New insights into pre-BCR and BCR signalling with relevance to B cell malignancies. Nature Reviews Immunology, 13(8):578–591, August 2013. ISSN 1474-1741. doi:10.1038/nri3487.

37. Farrell, M.V., Webster, S., Gaus, K. and Goyette, J. T cell membrane heterogeneity aids antigen recognition and t cell activation. Frontiers in Cell and Developmental Biology, 8, July 2020. doi:10.3389/fcell.2020.00609.

38. Ruthardt, N. and Bräuchle, C. Visualizing uptake and intracellular trafficking of gene carriers by single-particle tracking. In Topics in Current Chemistry, pages 283–304. Springer Berlin Heidelberg, 2010. doi:10.1007/128_2010_66.

39. Ponjavic, A., McColl, J., Carr, A.R., Santos, A.M., Kulenkampff, K., Lippert, A., Davis, S.J., Klenerman, D. and Lee, S.F. Single-Molecule Light-Sheet Imaging of Suspended T Cells. Biophysical Journal, 114(9):2200–2211, May 2018. ISSN 00063495. doi:10.1016/j.bpj.2018.02.044.

40. Wang, I.H., Burckhardt, C., Yakimovich, A. and Greber, U. Imaging, tracking and computational analyses of virus entry and egress with the cytoskeleton. Viruses, 10(4):166, March 2018. doi:10.3390/v10040166.

41. Dupont, A., Gorelashvili, M., Schüller, V., Wehnekamp, F., Arcizet, D., Katayama, Y., Lamb, D.C. and Heinrich, D. Three-dimensional single-particle tracking in live cells: news from the third dimension. New Journal of Physics, 15(7):075008, July 2013. doi:10.1088/1367-2630/15/7/075008.

42. Hou, S., Exell, J. and Welsher, K. Real-time 3d single molecule tracking. Nature Communications, 11(1), July 2020. doi:10.1038/s41467-020-17444-6.

43. Tolar, P., Hanna, J., Krueger, P.D. and Pierce, S.K. The Constant Region of the Membrane Immunoglobulin Mediates B Cell-Receptor Clustering and Signaling in Response to Membrane Antigens. Immunity, 30(1):44–55, January 2009. ISSN 1074-7613. doi:10.1016/j.immuni.2008.11.007.

44. Sohn, H.W., Tolar, P. and Pierce, S.K. Membrane heterogeneities in the formation of B cell receptor–Lyn kinase microclusters and the immune synapse. The Journal of Cell Biology, 182(2):367–379, July 2008. ISSN 0021-9525. doi:10.1083/jcb.200802007.

45. Whelan, D.R. and Bell, T.D.M. Image artifacts in Single Molecule Localization Microscopy: why optimization of sample preparation protocols matters. Scientific Reports, 5(1):7924, January 2015. ISSN 2045-2322. doi:10.1038/srep07924.

46. Beheiry, M.E. and Dahan, M. ViSP: representing single-particle localizations in three dimensions. Nature Methods, 10(8):689–690, August 2013. ISSN 1548-7091, 1548-7105. doi:10.1038/nmeth.2566.

47. Montiel, D., Cang, H. and Yang, H. Quantitative Characterization of Changes in Dynamical Behavior for Single-Particle Tracking Studies. The Journal of Physical Chemistry B, 110 (40):19763–19770, October 2006. ISSN 1520-6106. doi:10.1021/jp062024j. Publisher: American Chemical Society.

48. Aristov, A., Lelandais, B., Rensen, E. and Zimmer, C. ZOLA-3d allows flexible 3d localization microscopy over an adjustable axial range. Nature Communications, 9(1), June 2018. doi:10.1038/s41467-018-04709-4.

