## Supplementary Information for "High-density volumetric super-resolution microscopy"

### Supplementary Information:
### High-density volumetric super-resolution microscopy

**Sam Daly[a], João Ferreira Fernandes[b], Ezra Bruggeman[a], Anoushka Handa[a], Ruby Peters[c], Sarah Benaissa[d], Boya Zhang[d], Joseph S. Beckwith[a], Edward W. Sanders[a], Ruth R. Sims[e], David Klenerman[a], Simon J. Davis[b], Kevin O'Holleran[d], and Steven F. Lee[a,✉]**

[a]Yusuf Hamied Department of Chemistry, Lensfield Road, University of Cambridge, Cambridge, CB2 1EW, UK
[b]Radcliffe Department of Medicine and MRC Human Immunology Unit, John Radcliffe Hospital, University of Oxford, Oxford, OX3 9DU, UK
[c]Department of Physiology, Development and Neuroscience, University of Cambridge, Cambridge, CB2 3EL, UK
[d]Cambridge Advanced Imaging Centre, Downing Site, University of Cambridge, Cambridge, CB2 3DY, UK
[e]Wavefront-Engineering Microscopy Group, Photonics Department, Institut de la Vision, Sorbonne Université, INSERM, CNRS, Institut de la Vision, Paris, France

#### Contents

**S1 Single Molecule Light Field Microscopy** **S3**
S1.1 Optical setup . . . . . S3
S1.2 SMLFM design parameters . . . . . S3
S1.3 Improving contrast in SMLFM . . . . . S4

**S2 Realistic 3D-SMLM Simulations** **S5**
S2.1 Engineered PSF simulations . . . . . S5
S2.2 Light field simulations . . . . . S5
S2.3 Detection noise model . . . . . S6
S2.4 Simulation parameters . . . . . S6

**S3 Cramér–Rao lower bound** **S7**

**S4 Analysis of simulated data** **S8**
S4.1 Defining density . . . . . S8
S4.2 Defining speed improvement . . . . . S8

#### Supplementary Note S1: Single Molecule Light Field Microscopy

##### S1.1 Optical setup

The visual representation of the single molecule light field microscopy (SMLFM) platform in Fig. 1 is intentionally conceptual and reduced to key components. Here we present the optical platform in more detail. The optical set-up is based on a Fourier light field microscope, in which a microlens array (MLA) locally apertures the wavefront and generates an image at the detector (Fig. S1a). (1, 2) This results in a 2D image where the 3D position of single-molecule fluorescence is encoded via parallax (Fig. S1b). The hexagonal MLA (photographed in Fig. S1c) was specifically designed to fit within the conjugate back focal plane (BFP, Fig. S1d) to prevent partial illumination of microlenses. Partial lenslet illumination leads to distortions, such that the PSF could no longer be approximated by a 2D Gaussian function. (1) The hexagonal MLA fits in the BFP in its entirety and therefore all perspective views can be approximated by a 2D Gaussian function. As a result, the position of a point emitter in each perspective view is displaced in proportion to the average gradient of the wavefront (Fig. S1e), resulting in a point spread function comprising a hexagonal array of spots over an 8 μm axial range (Fig. S1f).

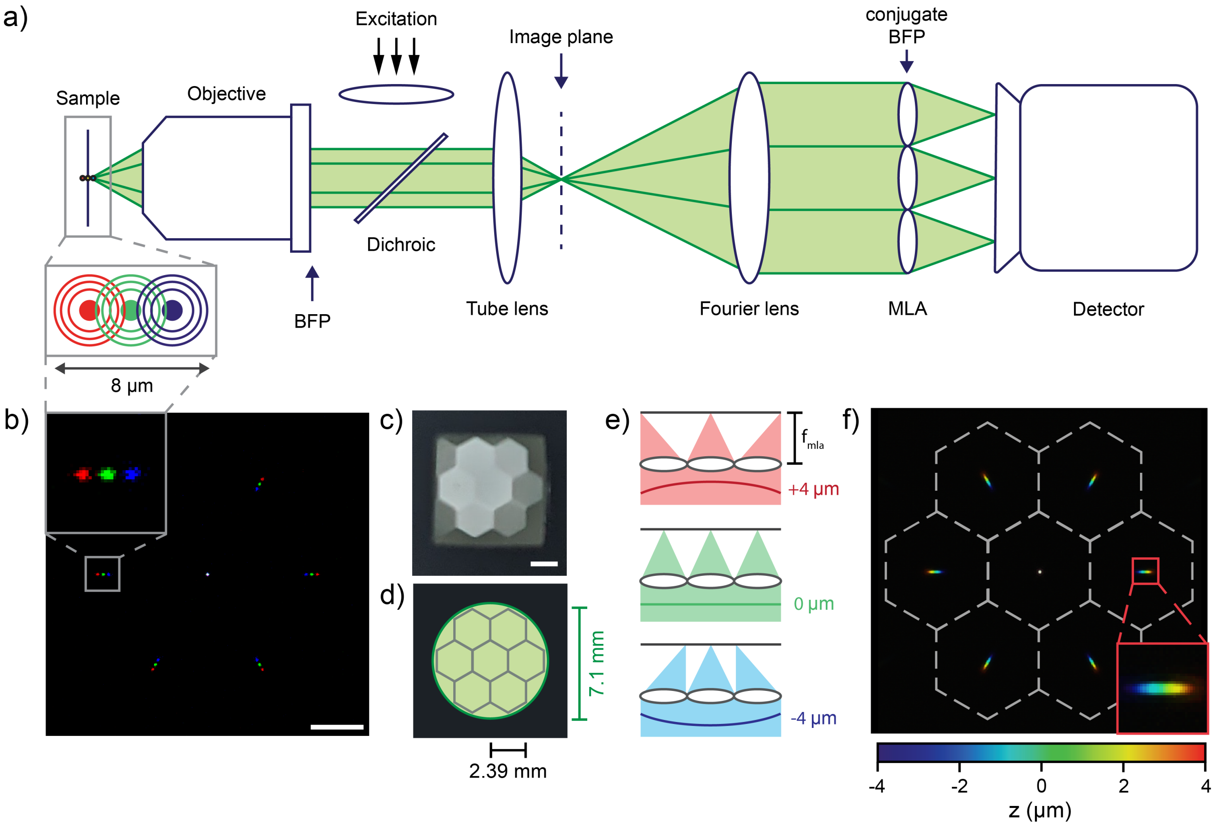


**Fig. S1. The single molecule light field microscopy (SMLFM) platform used in this study. a)** The optical layout showing a microlens array (MLA) segmenting the back focal plane (BFP) resulting in an image of the sample volume **b)** observed from 7 different perspective views. Scale bar is 20 μm. **c)** Photograph of the hexagonal MLA and **d)** the relative size of the MLA and the BFP. **e)** The MLA samples the curvature of the wavefront and displaces the focused spot in the outer perspective views (disparity) in proportion to the depth of the emitter. **f)** The point spread function (simulated) of the SMLFM platform presented herein over an 8 μm depth of field.

##### S1.2 SMLFM design parameters

Though consisting of few optical components, a number of design parameters govern the performance of FLFM, a detailed discussion of which can be found in (3). The number of illuminated microlenses along the diameter of the BFP, $N$, can be determined determined using Eq. S1. This is proportional to the number of perspective views of the sample volume.

$$N = \frac{d_{\mathrm{BFP}}}{\mathrm{P}} \tag{S1}$$

Where $d_{\text{BFP}}$ is the diameter of the BFP and P is the pitch of the MLA. In the present work, $N$ was experimentally measured to be $3.01$ by directly imaging the BFP and therefore 3 microlenses fit along the diameter of the BFP. The proportion of the BFP that is segmented by the MLA, also known as *occupancy ratio*, was therefore calculated to be $85\%$. In SMLFM, depth-of-field (DoF) and precision are limited by the low photon numbers generally yielded by single molecules. Therefore, the hexagonal MLA was implemented for the first time in this study in an attempt to improve photon throughput on previous SMLFM iterations that employed square MLAs.

The field-of-view (FoV) must match the spatial scales of the samples being imaged. In theory, the maximum theoretical FoV in SMLFM is that of the objective lens, however for simplicity we define a functional FoV that is the relative size of the central microlens in the BFP. This ensures that microlenses do not overlap. As such, the FoV can be deduced using Eq. S2.

$$\text{FoV} = \frac{\text{P}}{M} \qquad M = \frac{f_{\text{MLA}}}{f_{\text{objective}}}\frac{f_1}{f_2} \tag{S2}$$

Here, $M$ is the magnification of the optical system. The functional FoV obtained in the present work was calculated and measured as a circle of diameter 40 µm.

Another quantity which can be calculated is the maximum resolvability (R) of two point sources in each perspective view. This was determined to be 1.3 µm for this SMLFM setup and is described by Eq. S3.

$$\text{R} = N\frac{N}{2n_a} + 2\frac{\delta}{M} \tag{S3}$$

Where $\delta$ is the physical pixel size of the detector and $n_a$ is the objective numerical aperture. Finally, the DoF (which scales with the number of illuminated microlenses and hence the diameter of the BFP of the microscope) was determined experimentally to be ~8 µm and calculated at 7.8 µm using Eq. S4.

$$\text{DoF} = 2\lambda\frac{N^2}{n_a^2} + 2\frac{\delta}{M}\frac{N}{n_a} \tag{S4}$$

A summary of these parameters calculated for the SMLFM platform described in this work can be found in Table S1.

**Table S1.** Summary of the optical parameters of the SMLFM platform described in this study.

| Parameter (units) | Calculated or measured value |
|---|---|
| Number of illuminated microlenses, $N$ | 3.0 |
| Occupancy ratio (%) | 85 |
| Field-of-view (µm) | 40 × 40 |
| Resolvability, R (µm) | 1.3 |
| Depth-of-field (µm) | 7.8 |

##### S1.3 Improving contrast in SMLFM

An additional functional advantage of SMLFM is that unwanted fluorescence (originating from the solvent, coverslip, excitation beam, *etc.*) can be spatially localised in the BFP into a single perspective view (see Fig. 3a [left perspective view]). This cannot be achieved with other 3D modalities examined here as it is a consequence of segmenting the BFP with discrete microlenses that offer spatially defined zones where background fluorescence is collected. We find that the contrast enhancement affords higher localization precision because of a sharp increase in signal-to-noise ratio, despite globally reducing the maximum number of views available for 3D localization.

#### Supplementary Note S2: Realistic 3D-SMLM Simulations

##### S2.1 Engineered PSF simulations

Simulations of single-molecule images using the standard, astigmatic, double helix and tetrapod PSF were generated using a scaled Fourier transform of the electric field in the back focal plane of the objective due to an isotropic emitter at position $\mathbf{r} = (x_i, y_i, z_i)$. The electric field at the back focal plane of the objective is given by

$$\mathbf{E}_{\text{bfp}}(\rho,\phi;\mathbf{r}) = A(\rho)\exp\left(i\Phi_{\text{xyz}}(\rho,\phi;\mathbf{r})\right) \quad \textbf{(S5)}$$

where $A(\rho) = 1/\left(1-(\text{NA}\rho/n)^2\right)^{1/4}$ is an apodization factor for aplanatic collimation (4), NA is the numerical aperture of the objective, $n$ is the refractive index of the immersion and sample medium, $(\rho,\phi)$ are normalised polar coordinates in the back focal plane ($\rho$ is normalised to be 1 at the edge of the pupil), and the phase of the electric field is dependent on the position of the emitter in space (5):

$$\Phi_{\text{xyz}}(\rho,\phi;\mathbf{r}) = k_0\text{NA}\left(x_i\rho\cos\phi + y_i\rho\sin\phi + z_i\sqrt{\frac{n}{\text{NA}} - \rho^2}\right) \quad \textbf{(S6)}$$

The electric field at the image plane is calculated as the scaled Fourier transform of the electric field in the back focal plane:

$$\mathbf{E}_{\text{img}} = \mathcal{F}\left\{\mathbf{E}_{\text{bfp}}(\rho,\phi;\mathbf{r})\exp\left(i\Phi_{\text{mask}}(\rho,\phi)\right)\right\} \quad \textbf{(S7)}$$

where $\Phi_{\text{mask}}$ is the phase modulation due to a phase mask. The phase masks for the astigmatic and tetrapod PSFs were calculated respectively as scaled primary vertical astigmatism, and a linear combination of primary and secondary vertical astigmatism:

$$\Phi_{\text{mask,astig}} = \frac{2\pi}{5}\rho^2\cos(2\phi)$$
$$\Phi_{\text{mask,tetrapod}} = \frac{\pi}{2}\rho^2\cos(2\phi) - \frac{13\pi}{20}(4\rho^4 - 3\rho^2)\cos(2\phi)$$

The phase mask for the double helix PSF was calculated using vectorial phase retrieval using an experimental calibration z-scan of fluorescent beads with a diameter of 100 nm. (6)
The intensity distribution at the image plane is then calculated from the electric field as:

$$I = \mathbf{E}_{\text{img}}^{\dagger}\mathbf{E}_{\text{img}} \quad \textbf{(S8)}$$

where † is the adjoint operator.

##### S2.2 Light field simulations

For single-molecule light field simulations, the electric field in the back focal plane of the objective due to an isotropic emitter is described by Eq. (S5). The image formation by a microlens array is modeled as described in reference (7). The phase of the electric field is modulated by a hexagonal microlens array, and then Fresnel propagated over a distance equal to the focal length of the microlens array (8) as follows:

$$\mathbf{E}_{\text{img}} = \mathcal{F}^{-1}\left\{\mathcal{F}\left\{\mathbf{E}_{\text{bfp}}\exp\left(i\Phi_{\text{MLA}}\right)\right\}\times\exp\left[i\pi\lambda f_{\text{MLA}}\sqrt{f_x^2+f_y^2}\right]\right\} \quad \textbf{(S9)}$$

where the exponential term is the Fresnel transfer function, $f_x$ and $f_y$ are the spatial frequencies in the image plane, and $\mathcal{F}\{\}$ and $\mathcal{F}^{-1}\{\}$ represent the Fourier and inverse Fourier transform respectively. The phase modulation due to the microlens array is given by:

$$\Phi_{\text{MLA}}(\rho,\phi) = \left[\text{hex}\left(x_{uv},y_{uv}\right)\exp\left(\frac{-ik}{2f_{\text{MLA}}}(x_{uv}^2+y_{uv}^2)\right)\right] * \text{comb}(\rho,\phi) \quad \textbf{(S10)}$$

where the term between square brackets describes the phase of a single hexagonal microlens and $(x_{uv}, y_{uv})$ are normalised coordinates in a single microlens. The first term between the square brackets is a hexagonal binary amplitude mask, and the exponential term is the phase transformation due to a thin lens (9) with a focal length

$f_{\text{MLA}}$. The phase modulation induced by the entire MLA is obtained by tiling the phase modulation due to a single microlens in a hexagonal pattern through convolution with a hexagonal 2D comb function that specifies the position of the centers of the microlenses. The intensity at the image plane is calculated from the electric field in the image plane using Eq. (S8).

##### S2.3 Detection noise model

Detection by an EMCCD camera is modelled as in (10)

$$n_{ie}(x,y) = \mathcal{P}\{\text{QE} \cdot [I(x,y) + b] + c\} \tag{S11}$$

$$I_{\text{camera}}(x,y) = \Gamma\{n_{ie}(x,y), g_{em}\} + \mathcal{G}(0, \sigma_{rn}) + \mathcal{O} \tag{S12}$$

where $I(x,y)$ is the intensity distribution at the image plane in photons, $b$ is a constant fluorescence background in photons, $n_{ie}(x,y)$ is the number of input electrons, $\mathcal{P}$ represents a Poisson distribution, QE is the quantum efficiency of the camera, $c$ is the spurious charge, $\Gamma$ represents a gamma function, $g_{em}$ is the electron multiplication gain, $\mathcal{G}$ represents a Gaussian distribution with a zero mean and standard deviation $\sigma_{rn}$, and $\mathcal{O}$ is a constant offset. Detector saturation is taken into account.

##### S2.4 Simulation parameters

Simulations were performed for standard, astigmatic, double helix, tetrapod and light field PSF at different emitter densities (between N$=2$ and N$=150$ emitters/volume), with three times 100 independent repeats for each density. A fixed number of emitters N were randomly placed in a 3D volume with a specific DoF and FoV as summarised in Table S2. The fluorescence background was fixed across simulations and set to 10 photons per 266 nm pixel. For the light field simulations, the background per pixel was divided by the number of microlenses in the pupil (seven in this case), as the background is also divided into the different perspective views. Simulations were performed for three signal levels: 1,000, 4,000, and 10,000 detected photons/emitter. For each PSF, a z-stack was also simulated of a single bright emitter (10,000 detected signal photons, no background photons) covering the full DoF of the method for calibration.

**Table S2.** Simulation parameters.

| PSF | DoF (µm) | FoV (µm$^2$) | Pixel size (nm) |
|---|---|---|---|
| Standard | 0 | $20 \times 20$ | 110 |
| Astigmatic | 1 | $20 \times 20$ | 110 |
| Double helix | 4 | $20 \times 20$ | 266 |
| Light field | 8 | $20 \times 20$ | 266 |
| Tetrapod | 8 | $20 \times 20$ | 110 |

The following system parameters were used for the light field simulations: hexagonal microlens array with a pitch of 2.39 mm, microlens focal length of 175 mm, fourier lens focal length of 175 mm, tube lens focal length of 200 mm (Nikon), and a 60$\times$ water-immersion objective (Nikon) with NA = 1.27. The camera parameters match the Evolve Delta 512 EMCCD camera from Photometrics, *i.e.* a physical pixel size of 16 µm.

The following values were used for the detection noise parameters were used: QE $=1$ electrons/photon, $c = 0.002$ electrons, $g_{em} = 250$ counts/electron (counts = analog-to-digital unit), $\sigma_{rn} = 74.4$ electrons, $\mathcal{O} = 400$ counts. These parameters correspond to the EMCCD camera Evolve Delta 512 from Photometrics.

#### Supplementary Note S3: Cramér–Rao lower bound

The Cramér-Rao lower bound (CRLB) is the theoretical limit of the variance of an unbiased estimator, in this case the 3D localization of a molecule. Assuming Poisson distributed noise, the CRLB can be determined by the inverse of the Fisher information matrix, as described in (11) and Eq. S13.

$$F_x = \sum_{k=1}^{N} \frac{1}{\mathrm{S}\,(\mathrm{k}) + b} \left( \frac{\partial \mathrm{S}\,(\mathrm{k})}{\partial x} \right)^2 \tag{S13}$$

Where N is the total number of pixels on the sensor, S(k) is the signal for the $k^{th}$ pixel and b is the background signal. The signal, S(k), is calculated as the PSF modeled as described in Supplementary Note S2 section S2.2 using parameters that match the experimental setup detailed in the Methods.

The hexagonal SMLFM platform described in the present work was simulated for 4,000 and 50,000 detected photons to determine the maximum theoretical precision of the instrument for typical experimental photon values and an unlimited photon scenario. The CRLB was computed for these images over the whole axial range (8 µm) and presented in Fig. S2.

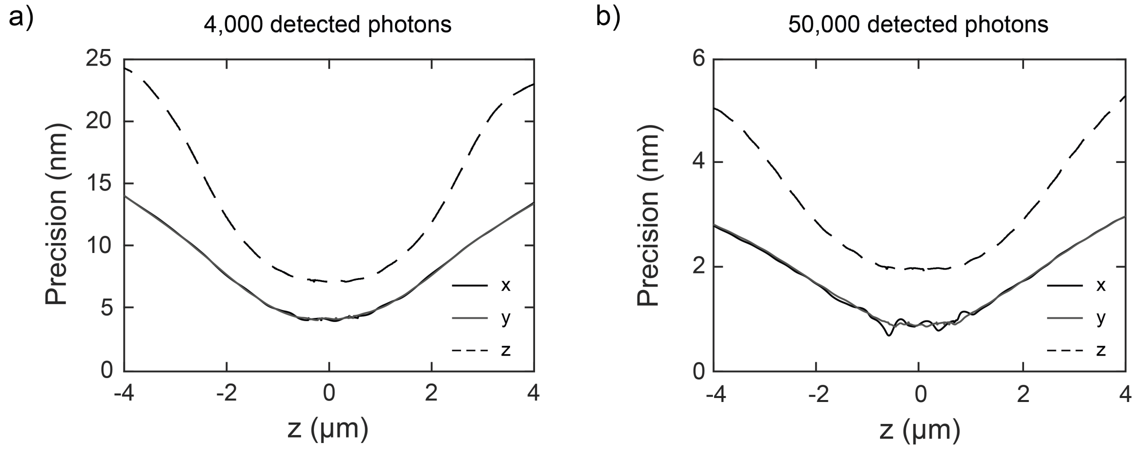


**Fig. S2. Lateral and axial Cramér–Rao lower bounds.** Lateral and axial Cramér–Rao lower bounds for the experimental setup discussed in this work determined through simulations for **a)** 4,000 detected photons and **b)** 50,000 detected photons with 150 background photons.

The precision of the optical setup at typical single-molecule photon values was determined to be below 14 nm laterally and 24.5 nm axially over the whole 8 µm DoF. Precision reaches a maximum of 5 nm laterally and 8nm axially at the middle of the focal volume for 4,000 photons. This improves to $\leq$2 nm in the photon-unlimited case. Although the lateral and axial precision is not isotropic across the whole DoF these results are consistent with previous work (1) that employed a square MLA and exhibits improved localisation precision throughout.

#### Supplementary Note S4: Analysis of simulated data

##### S4.1 Defining density

In this study we define *density of emitters* as the number of single emitters that give rise to an intensity distribution on a 2D detector per frame (per unit area). This approach aligns with previous work (12) and is consistent with the projection of a 3D volume onto a 2D array of pixels. A consistent $20 \times 20$ μm FoV meant that emitter densities spanned the range $0.005$ to $0.375$ $\mu m^{-2}$ with $0.100$ $\mu m^{-2}$ resembling a typical labelling density for 2D-SMLM experiments. (13) Fig. S3 shows the variation in the PSF size throughout the corresponding DoF established through intensity thresholds.

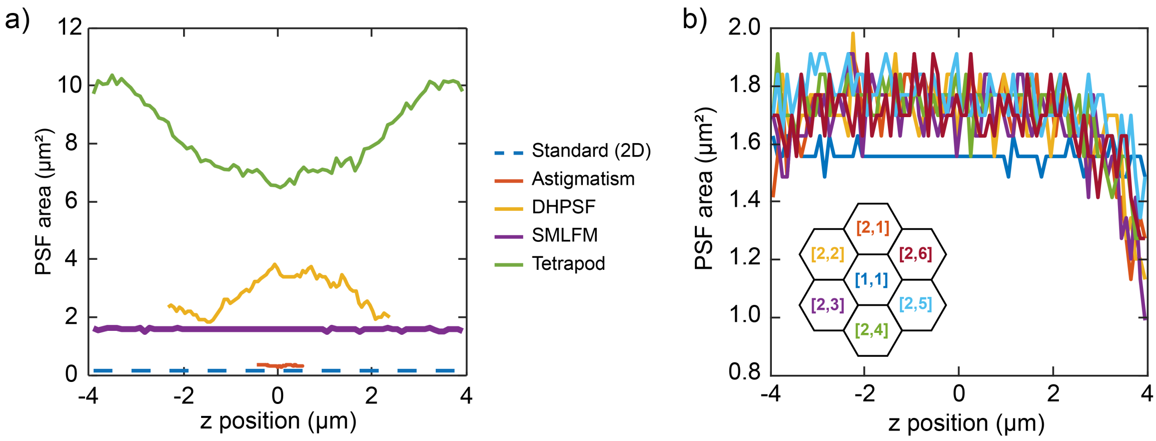


**Fig. S3. Relative size of 3D PSFs as a function of axial position. a)** Comparative area of 3D PSFs explored herein at different axial planes within the DoF determined by realistic calibration datasets using an intensity-based threshold. Note the light field PSF (central view) is size invariant across the whole DoF, a result of the low effective numerical aperture of each microlens and is what enables the SMLFM PSF to be fitted with conventional 2D Gaussian fitting prior to 3D reconstruction. **b)** SMLFM PSF size as a function of axial position for each perspective view determined by an intensity threshold. The axially invariant size of the outer perspective views behaves identically between perspective views and similarly to the central view which resides over only a marginally smaller detector area.

##### S4.2 Defining speed improvement

We quantified the maximum speed difference between SMLFM and the current state-of-the-art DHPSF (with which we possess most experimental experience.) When determining this speed enhancement it was important to consider the optimal localization densities for both techniques to ensure a fair comparison. Therefore, we compared the emitter density at which both techniques shared the same error rate (*i.e.* the value at which the proportion of true positive localizations relative to the ground truth was equal). Plots of error rate vs. density are presented in Fig. S4 with the difference between the two interpolated curves (corresponding to the relative speed of each technique) plotted directly above.

Speed improvement was also calculated at different values of detected photons to reflect different labelling scenarios: a fluorescent protein (1,000 photons), an organic dye (4,000 photons) and a next-generation probe (10,000 photons). We determined SMLFM to facilitate a maximum speed improvement of 25.3× (40.0% error rate), 8.95× (13.4% error rate) and 10.6× (13.4% error rate) at 1,000, 4,000 and 10,000 detected photons, respectively.

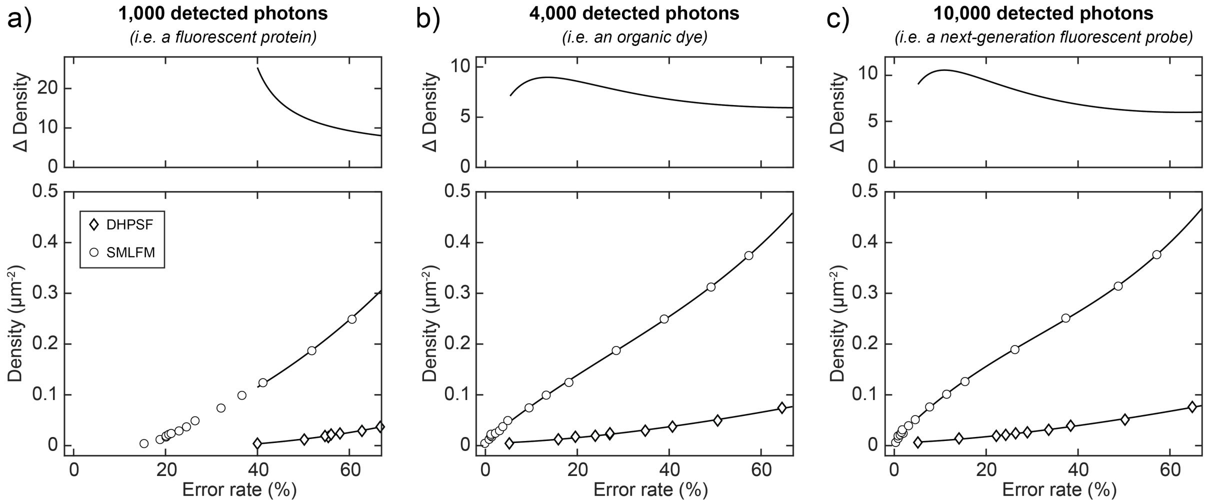


**Fig. S4. Relative speed of SMLFM compared to the DHPSF for typical photon values.** The density of single emitters per frame for a given error rate (proportion of false negatives relative to the number of ground truth localizations) for SMLFM and DHPSF simulated at **a)** 1,000 detected photons, **b)** 4,000 detected photons and **c)** 10,000 detected photons. Plots of the density difference between the two interpolated lines (relative speed) are presented above each graph. To avoid extrapolation, at 1,000 photons a density difference is only calculated after an error rate of 40%.

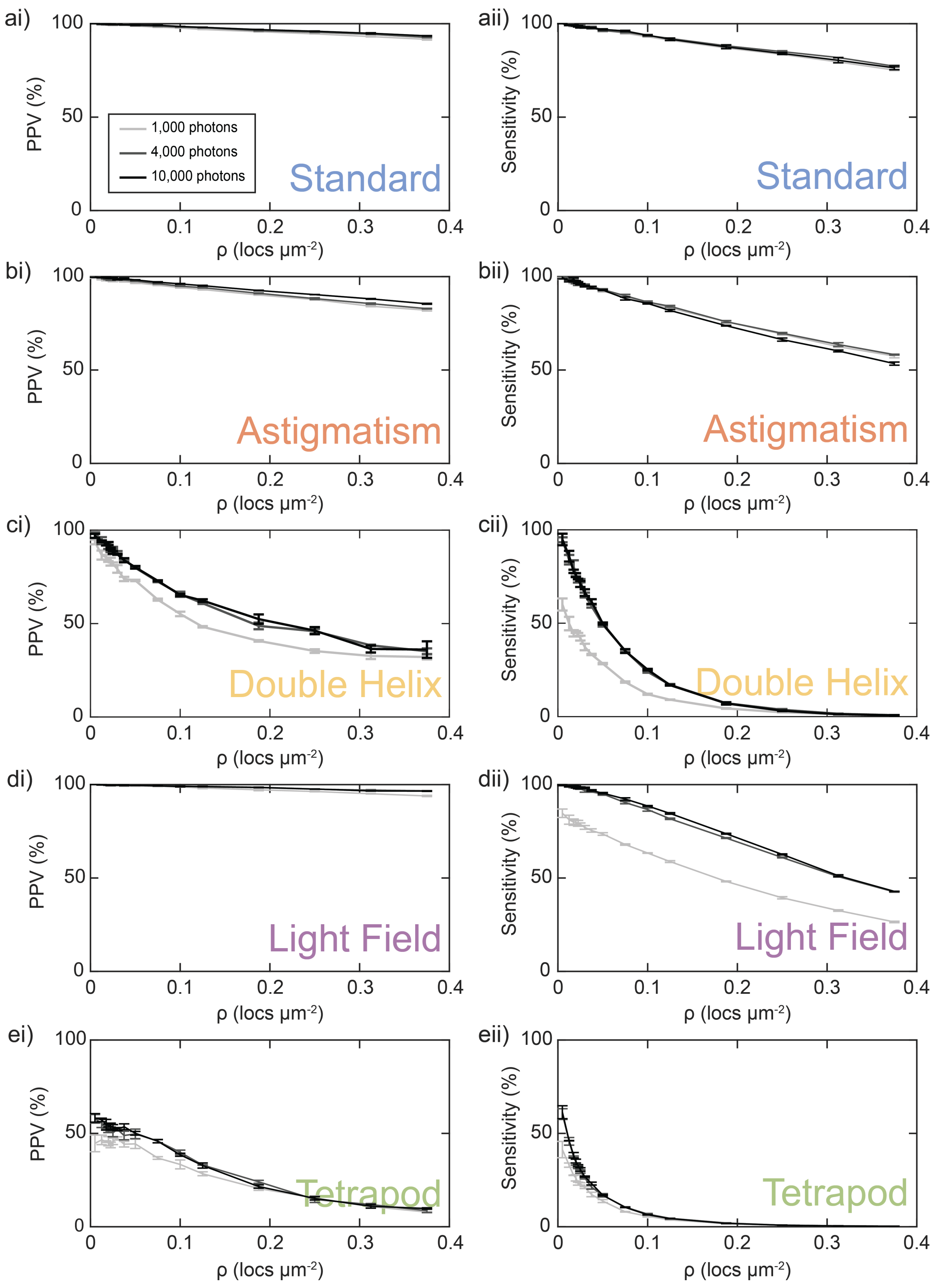


**Fig. S5. Positive predictive value (PPV) and sensitivity plots at different signal-to-noise ratios. a–ei)** PPV and **a–eii)** sensitivity plots for each PSF explored here, including the standard, astigmatic, double helix, light field and tetrapod PSFs simulated for 1,000, 4,000 and 10,000 detected signal photons.

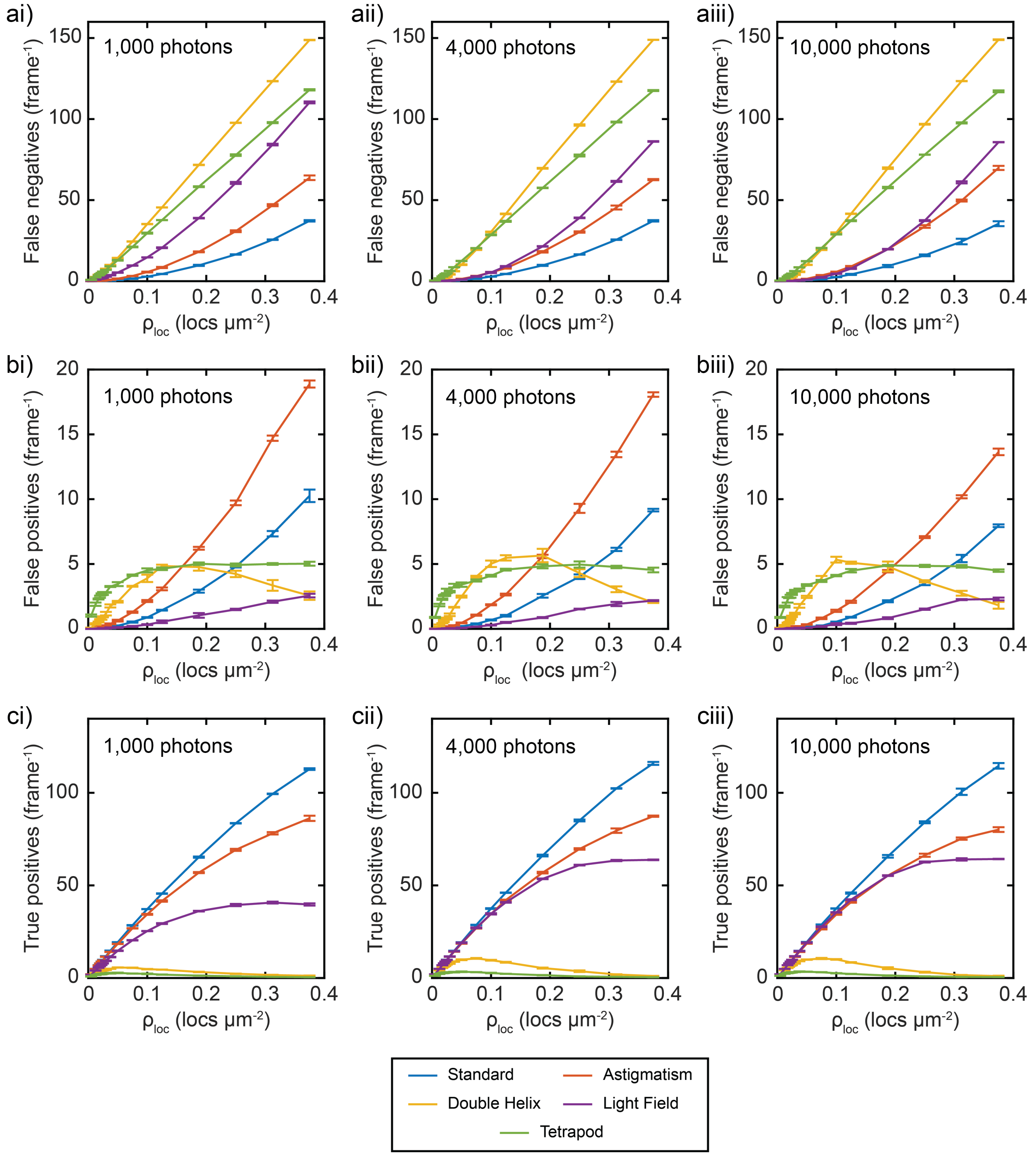


**Fig. S6. Imaging quality metrics from the analysis of simulated localization datasets. a)** Number of false negative localizations per frame for each SMLM technique simulated here-in at **i)** 1,000, **ii)** 4,000, **iii)**, and 10,000 detected photons determined through simulations described in Supplementary Note S2. **b)** Number of false positive localizations per frame at **i)** 1,000, **ii)** 4,000, **iii)**, and 10,000 detected photons. **c)** Number of true positive localizations per frame at **i)** 1,000, **ii)** 4,000, **iii)**, and 10,000 detected photons. Each plot represents the average value over three repeats where error bars represent one standard deviation. A rotation filter was applied to each frame of the matching analysis to account for the higher likelihood the false pairing of a fitted point to ground truth coordinate at higher localisation densities. This filter rotated the fitted data 90° relative to the ground truth coordinates per frame and if another localisation occurs within the mean square distance tolerance then this coordinate was discarded.

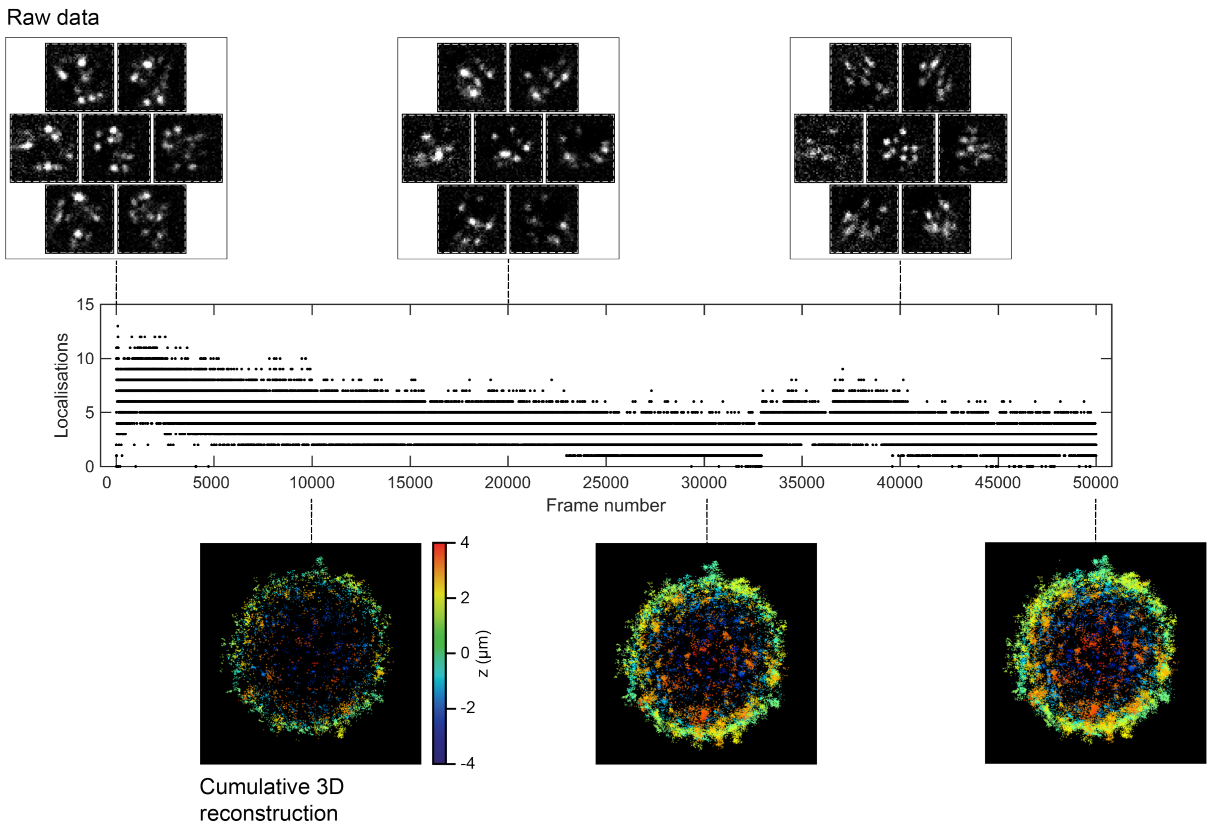


**Fig. S7. Localization rate during dSTORM imaging of the BCR.** Single cropped frames of raw data is presented on top and the cumulative 3D reconstruction below. An additional laser at 405 nm was used commencing at frame 33,000 and increased in power at frame 36,000 to ensure all fluorophores had been imaged.

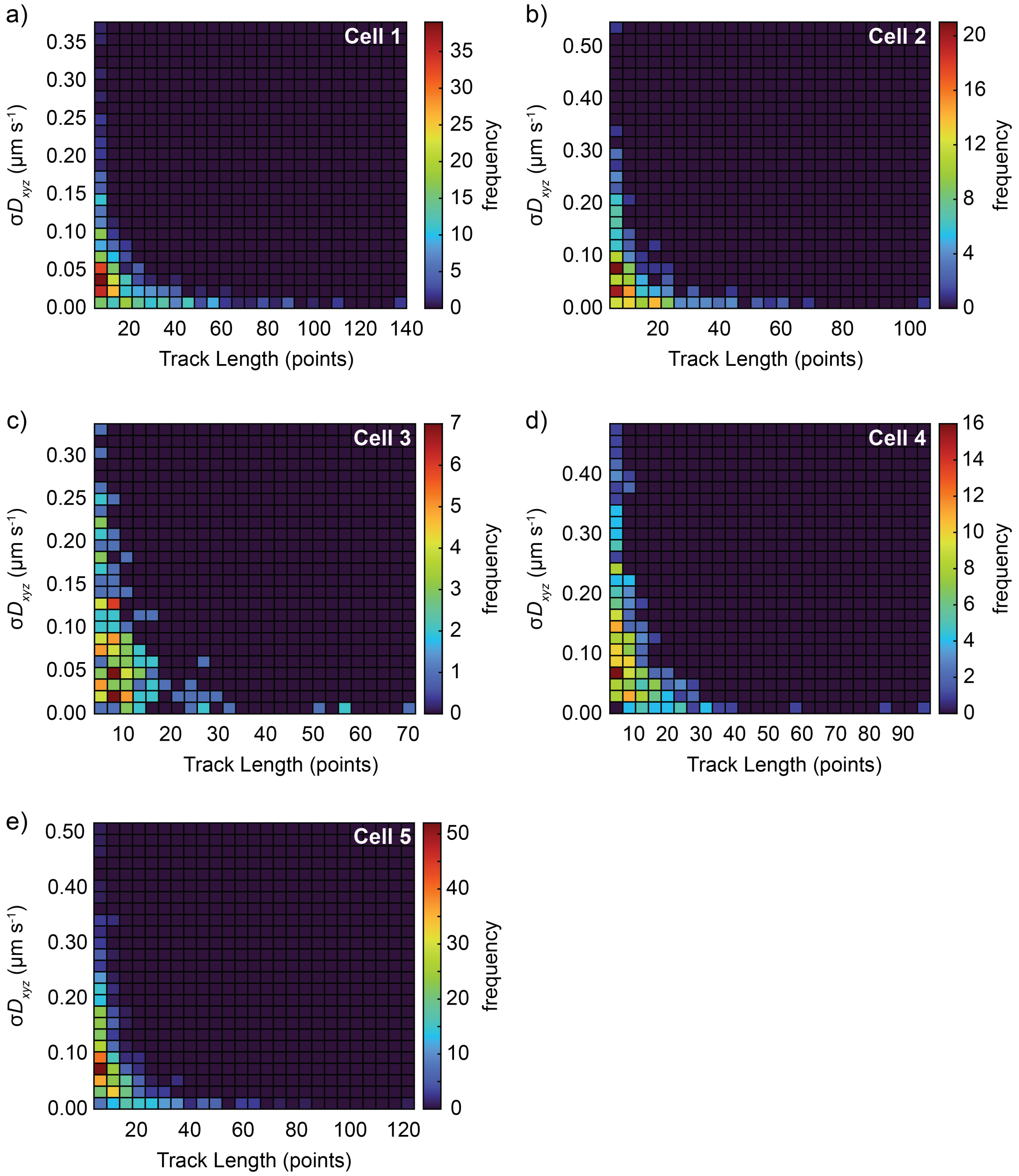


**Fig. S8. Maximum likelihood error for diffusion measurements.** 2D histograms showing the maximum likelihood error for diffusion measurements in xyz as a function of track length for each cell (**a**–**e**) that contributes to the data presented in Fig. 4.

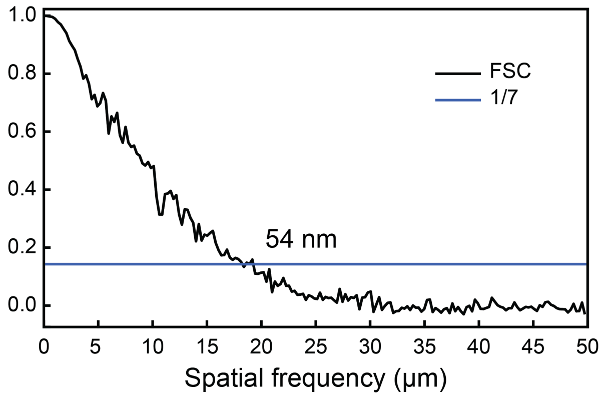


**Fig. S9. Resolution of high-density tubulin datasets.** Fourier shell correlation (FSC) estimates an isotropic resolution of 54 nm at a 1/7 cutoff for the high-density tubulin image presented in Fig. 5.

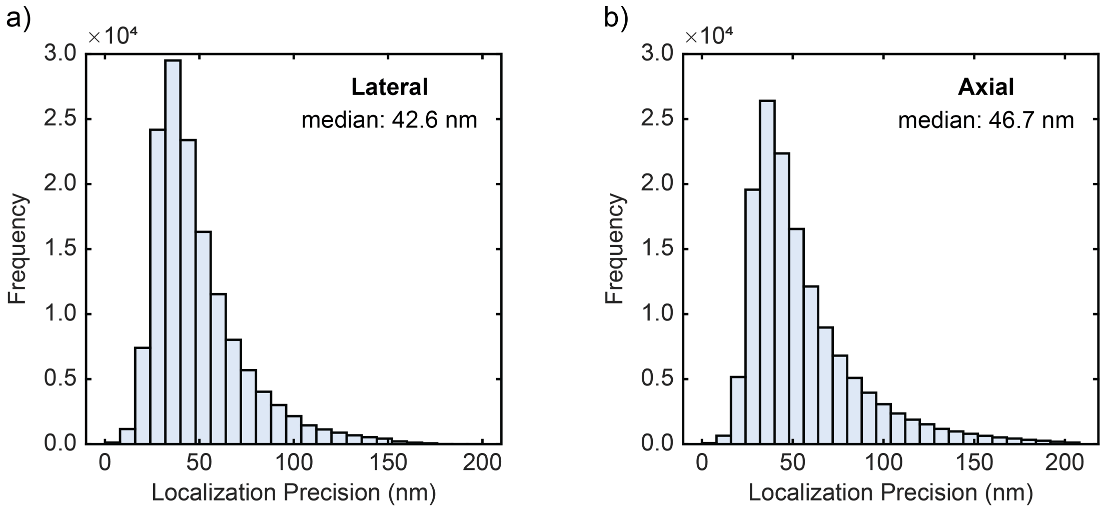


**Fig. S10. Lateral and axial localization precision in tubulin image.** Localization precision obtained **a)** laterally and **b)** axially in the high-density tubulin image presented in Fig. 5.

### References

1. Sims, R.R., Abdul Rehman, S., Lenz, M.O., Benaissa, S.I., Bruggeman, E., Clark, A., Sanders, E.W., Ponjavic, A., Muresan, L., Lee, S.F. and O'Holleran, K. Single molecule light field microscopy. *Optica*, 7(9):1065, September 2020. ISSN 2334-2536. doi: 10.1364/OPTICA.397172.
2. Guo, C., Guo, C., Liu, W., Liu, W., Hua, X., Li, H. and Jia, S. Fourier light-field microscopy. *Optics Express*, 27(18):25573–25594, September 2019. ISSN 1094-4087. doi: 10.1364/OE.27.025573. Publisher: Optical Society of America.
3. Galdón, L., Saavedra, G., Garcia-Sucerquia, J., Martínez-Corral, M. and Sánchez-Ortiga, E. Fourier lightfield microscopy: a practical design guide. *Applied Optics*, 61(10):2558, March 2022. doi: 10.1364/ao.453723.
4. Kim, J., Wang, Y. and Zhang, X. Calculation of vectorial diffraction in optical systems. *JOSA A*, 35:526–535, 2018.
5. Botcherby, E., Juskaitis, R., Booth, M. and Wilson, T. An optical technique for remote focusing in microscopy. *Optics Communications*, 281:880–887, 2008.
6. Ferdman, B., Nehme, E., Weiss, L.E., Orange, R., Alalouf, O. and Shechtman, Y. VIPR: vectorial implementation of phase retrieval for fast and accurate microscopic pixel-wise pupil estimation. *Optics Express*, 28(7):10179–10198, March 2020. ISSN 1094-4087. doi: 10.1364/OE.388248. Publisher: Optica Publishing Group.
7. Guo, C., Liu, W., Hua, X., Li, H. and Jia, S. Fourier light-field microscopy. *Optics Express*, 27(18):25573–25594, 2019.
8. Voelz, D. *Computational fourier optics : a MATLAB tutorial*. SPIE, 1st edition, 2011.
9. Goodman, J. *Introduction to Fourier Optics*. W.H.Freeman, Co Ltd, 3rd edition, 2005.
10. Sage, D., Pham, T.A., Babcock, H., Lukes, T., Pengo, T., Chao, J., Velmurugan, R., Herbert, A., Agrawal, A., Colabrese, S., Wheeler, A., Archetti, A., Rieger, B., Ober, R., Hagen, G.M., Sibarita, J.B., Ries, J., Henriques, R., Unser, M. and Holden, S. Super-resolution fight club: assessment of 2D and 3D single-molecule localization microscopy software. *Nature Methods*, 16(5):387–395, May 2019. ISSN 1548-7105.
11. Ober, R.J., Ram, S. and Ward, E.S. Localization accuracy in single-molecule microscopy. *Biophysical Journal*, 86(2):1185–1200, February 2004. doi: 10.1016/s0006-3495(04)74193-4.
12. Nehme, E., Freedman, D., Gordon, R., Ferdman, B., Weiss, L.E., Alalouf, O., Naor, T., Orange, R., Michaeli, T. and Shechtman, Y. DeepSTORM3d: dense 3d localization microscopy and PSF design by deep learning. *Nature Methods*, 17(7):734–740, June 2020. doi: 10.1038/s41592-020-0853-5.
13. van de Linde, S., Löschberger, A., Klein, T., Heidbreder, M., Wolter, S., Heilemann, M. and Sauer, M. Direct stochastic optical reconstruction microscopy with standard fluorescent probes. *Nature Protocols*, 6(7):991–1009, June 2011. doi: 10.1038/nprot.2011.336.
